# Chronic nicotine increases midbrain dopamine neuron activity and biases individual strategies towards reduced exploration in a foraging task

**DOI:** 10.1101/2021.01.29.428835

**Authors:** Malou Dongelmans, Romain Durand-de Cuttoli, Claire Nguyen, Maxime Come, Etienne K. Duranté, Damien Lemoine, Raphael Britto, Tarek Ahmed Yahia, Sarah Mondoloni, Steve Didienne, Elise Bousseyrol, Bernadette Hannesse, Lauren M. Reynolds, Nicolas Torquet, Deniz Dalkara, Fabio Marti, Alexandre Mourot, Jérémie Naudé, Philippe Faure

## Abstract

Long-term exposure to nicotine alters brain circuits and induces profound changes in decision-making strategies, affecting behaviors both related and unrelated to drug seeking and consumption. Using an intracranial self-stimulation reward-based foraging task, we investigated the impact of chronic nicotine on the trade-off between exploitation and exploration, and the role of ventral tegmental area (VTA) dopamine (DA) neuron activity in decision-making unrelated to nicotine-seeking. Model-based and archetypal analysis revealed a substantial inter-individual variability in decision-making strategies, with mice passively exposed to chronic nicotine visiting more frequently options associated with higher reward probability and therefore shifting toward a more exploitative profile compared to non-exposed animals. We then mimicked the effect of chronic nicotine on the tonic activity of VTA DA neurons using optogenetics, and found that photo-stimulated mice had a behavioral phenotype very close to that of mice exposed to nicotine, suggesting that the dopaminergic control of the exploration/exploitation balance is altered under nicotine exposure. Our results thus reveal a key role of tonic midbrain DA in the exploration/exploitation trade-off and highlight a potential mechanism by which nicotine affects decision-making.

## Introduction

Nicotine is the primary reinforcing component driving tobacco addiction ^1,2 3^. Like most addictive substances, nicotine is hypothesized to perpetuate addiction through alterations in dopamine (DA) signaling and plasticity in the mesocorticolimbic pathway ^4^. Repeated activation of ventral tegmental area (VTA) DA neurons by nicotine not only leads to reinforcement but also to craving and lack of self-control over intake ^5^. Concurrently, chronic exposure to nicotine also causes modifications of decision-making processes, which affect personality traits and behaviors that extend beyond drug-seeking or -consumption ^6,7^, such as impulsivity ^8,9^ and exploratory behaviors ^10,11^. These traits in turn actively contribute to the persistence of drug consumption, by promoting relapse and susceptibility to other addictions ^12^. However, the impact of nicotine-induced modifications of DA neural networks on choice behaviors, and particularly the tradeoff between exploration and exploitation, is still undetermined.

When faced with a choice between two alternatives with low and high probabilities of reward, animals choose the less likely rewarded option a significant portion of the time. The origin of such seemingly suboptimal choice strategy remains poorly understood. It has been interpreted in different studies as noise, error, risk seeking, irrational belief or exploration ^7,13–16^. In the context of exploration, choosing an option with less likelihood of immediate reward is an essential adaptive process related to cognitive flexibility and to gathering information about unknown or uncertain outcomes in a changing environment. Exploration is thus central to the emergence and organization of behaviors ^17^, naturally resulting in the acquisition of new information crucial for learning and optimizing behavioral strategies ^7,13^. Determining whether chronic nicotine exposure alters such exploratory behaviors is thus fundamental to help understand modifications of individual traits associated with continued nicotine consumption.

Altered DA function is a promising candidate to link chronic nicotine exposure to changes in decision making behavior. This neuromodulator, which is at the crossroads of motivation, learning and decision-making, can be hijacked, in the context of addiction, by most drugs of abuse ^18–20^. Changes in the spontaneous tonic firing of VTA DA neurons, as a consequence of repetitive drug-use, can indeed alter the subjective value assigned to available rewards ^19^, as well as the motivational salience of the drug or of drug-predicting cues ^21^, influencing decisions about which reward to pursue ^22^. Tonic DA can scale the performance of a learned behavior ^23^, the incentive value associated with environmental stimuli ^24^, or signal the average reward ^25^. In the exploration/exploitation framework, the role of tonic DA remains debated. The effect of DA manipulation on the exploration/exploitation balance is convincing but varies depending on the task ^26–28^. Increasing tonic striatal DA release has been suggested to either increase ^28^ or decrease ^27^ the level of exploration. Decreasing tonic striatal DA has also been suggested to increase exploration ^29^. Hence, drug-induced alterations of DA transmission may modify behavioral choices, either positively or negatively depending on the environment and the specific type of DA manipulation.

Using an intracranial self-stimulation (ICSS) reward-based foraging task for mice, we have shown that decisions in this foraging task are modulated by the cholinergic neurotransmission of the VTA, with a particular role of nicotinic acetylcholine receptors (nAChR) in expected uncertainty driven exploration ^30^. Here we demonstrated that chronic nicotine exposure increases the tonic activity of VTA DA neurons and reduces exploration, with mice focusing on the most valuable options at the expense of information gathering. Increasing the tonic activity of VTA DA neurons using optogenetics was sufficient to mimic the behavioral bias (or exploratory decrease) induced by nicotine, indicating that the DA control of the exploration/exploitation balance is altered by long-term nicotine exposure.

## Results

### Mice biased their choices in a multi-armed ICSS bandit task by motor cost, probability and uncertainty of the reward delivery

To assess choice behavior in an uncertain environment, we used a multi-armed ICSS bandit task for mice where specific locations, hereafter called targets, were associated with brain stimulation rewards delivered to the medial forebrain bundle (MFB) (Figure 1A, Supplementary Figure 1) ^16,30,31^. The task takes place in a circular open-field (interior diameter = 80 cm), with three explicitly marked targets forming the apices of a triangle (Figure 1B). Passing over each target results in the delivery of a rewarding intra-cranial electrical stimulation. Mice could not receive two consecutive stimulations from the same target, and thus learn to forage from one to another to continue receiving stimulations (Figure 1B Left). During the training period (5-min daily sessions), hereafter called the deterministic setting (DS, Figure 1C Left), every visit to a target was reinforced by a stimulation reward (reward probability p = 100% at each location, p_100_). At the end of the DS, mice were confronted with a probabilistic setting (PS, Figure 1C Right) where each target was now associated with a different probability of stimulation delivery (p = 100%, 50% and 25%, Figure 1C Right). As previously shown ^30^, both the expected reward probabilities and uncertainties associated with the different targets in the PS induced a marked change in the behavioral pattern compared to the DS. Trajectories at the end of the DS were stereotyped, almost circular, with a low probability of directional changes (i.e returning to the previous target, Figure 1D) due to an associated motor cost ^16^. In contrast, in the PS mice distributed their choices differently and increased their probability of directional changes, indicating an adaptation from the circular strategy (Figure 1D). Directional changes in the PS were not random: rather, they allowed animals to focus on specific targets. Indeed, compared to the DS where mice visited the three targets with a uniform distribution, in the PS mice visited more often the targets associated with the highest reward probabilities (i.e. p_100_ and p_50_, Figure 1E). This indicates a matching behavior, i.e., a quantitative relationship between the rate of target visits and the probability to receive a reward on each target. Since mice could not receive two consecutive rewards from the same target, this repartition on the rewarding locations resulted from a sequence of binary choices (Figure 1F) in three gambles (G_100_, G_25_, G_50_) between two respective payoffs (here, G_100_ = {p_50_ versus p_25_}, G_25_ = {p_100_ versus p_50_}, G_50_ = {p_100_ versus p_25_}). Hence, we analyzed the sequence of choice statistics using a transition function, allowing us to investigate each gamble independently. For G_100_ and G_50_, mice chose the optimal location (i.e., the one associated with the highest probability of reward) more than 50% of the time. However, as previously observed, for G_25_ the probability to choose p_100_ over p_50_ was not different from a random choice (Figure 1F), which has been interpreted as indicating that mice assign a positive motivational value to expected uncertainty, which is maximal at p_50_ ^30^. Overall mice biased their choices depending on both the probability and the uncertainty of reward delivery. Behavior in the task was therefore the result of a combination between rewards, uncertainty and motor cost.

**Figure 1:**
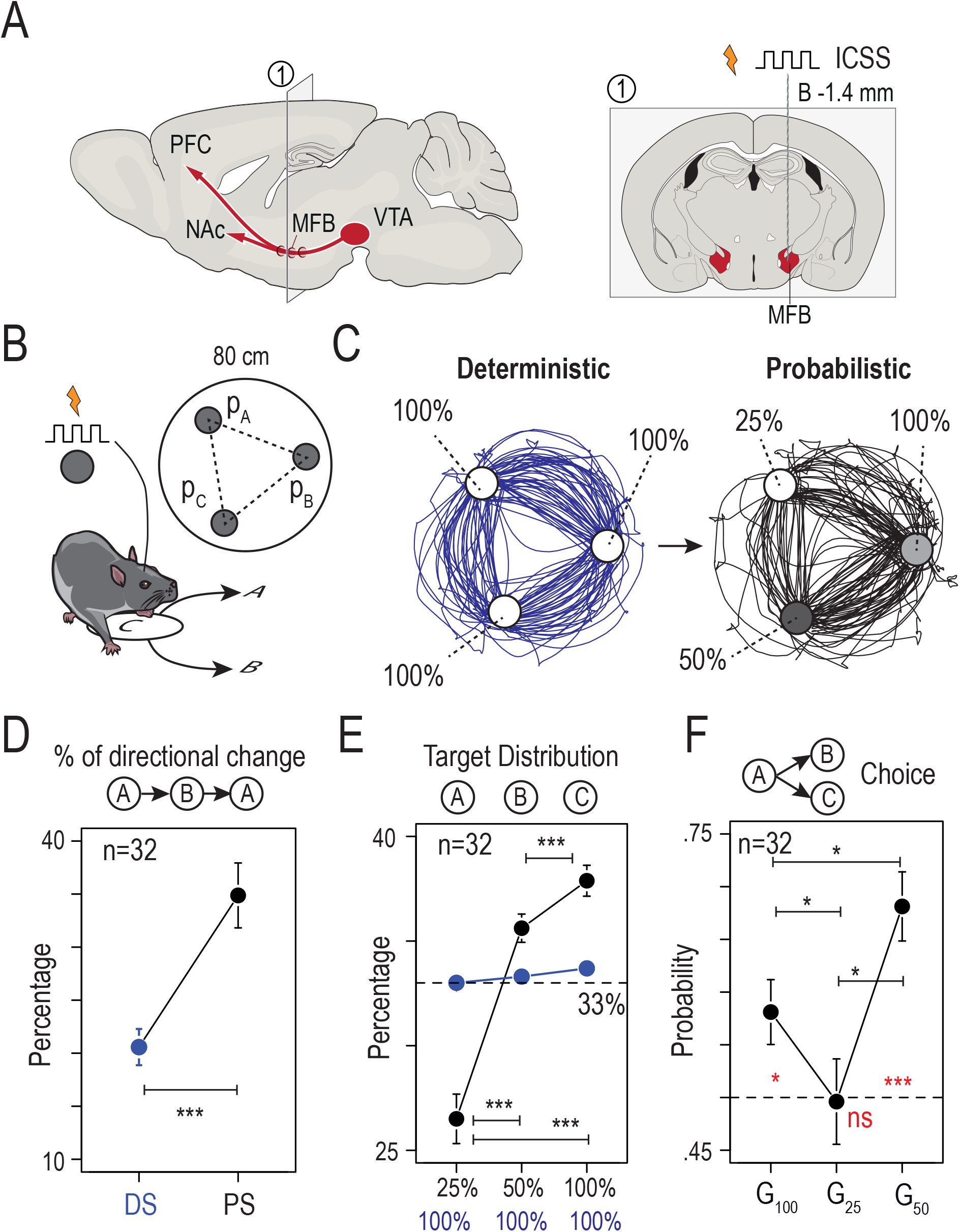
Mice exhibited suboptimal behavior and exploratory choices in a spatial version of a multi-armed bandit task with probabilistic settings. (A) Mice were implanted unilaterally with bipolar stimulation electrodes to deliver electrical stimulation at the level of the medial forebrain bundle in order to support intracranial self-stimulation (ICSS) behavior. *Right*: A coronal section of the mouse brain illustrating a representative electrode positioned in the MFB at Bregma −1.4 mm AP. (B) Schematic of the behavioral paradigm: mice are placed in a circular open-field (interior diameter = 80 cm), with three equidistant targets (A, B, and C - labelled on the open field floor) that are associated with a given probability (P_A_, P_B_, _or_ P_C_) of ICSS reward delivery when the animal is detected in a 60 mm zone around the target. (C) Sample trajectories for one mouse under the deterministic setting (DS) of the task, in which each of the three targets were rewarded by an ICSS with P = 100% (left panel, blue), and in the probabilistic setting (PS), in which the three targets were associated with distinct probabilities of ICSS delivery (P_A_ = 100, P_B_ = 50 and P_C_ = 25 %) (right panel, black). Two stimulations could not be delivered consecutively in the same zone, therefore animals learned to alternate between targets with a circular pattern in the DS (blue), and a less stereotyped pattern in the PS (black). (D) Comparison of the percentage of directional changes during DS (blue) and PS (black) (Wilcoxon signed rank test, p < 0.001, n = 33). (E) Repartition of visits to the three targets. Under the DS (blue), animals distributed uniformly their choices of visiting each of the three options (around 33%, Friedman rank sum test, p = 0.82). During the PS (black), animals reorganized their behavior and visited more frequently options with greater probabilities of reward (Friedman rank sum test, p < 0.001, and paired Wilcox Test p < 0.001 for the three comparisons, n = 33). (F) Probability to choose the option with the highest probability of reward for the three possible gambles: G_100_ = choice of 50% over 25%, G_25_ = choice of 100% over 50 and G_50_ = choice of 100% over 25%. Red asterisk: Comparison with a true mean of 0.5 (One Sample t-test with Holm correction, n = 33) for G_100_ (p = 0.026), G_25_ (p = 0.92) and G_50_ (p < 0.001). Black asterisk: Paired comparison (Paired t-test with Holm correction, n = 33) for G_100_-G_25_ (p = 0.03), G_100_-G_50_ (p = 0.048) and G_25_-G_50_ (p = 0.025).

### Chronic nicotine exposure decreased exploration and increased exploitation of the most valuable options

We aimed to investigate the effects of chronic nicotine exposure on decision-making behavior and on the balance between exploration and exploitation. To do so, we implanted osmotic minipumps subcutaneously to expose mice to continuous nicotine (Nic, 10mg/kg/day) or saline (Sal) for 3 weeks and then compared their behavior in the PS of the ICSS task (Figure 2A). Because nicotine induces long lasting adaptations in the midbrain DA system ^32^, and because VTA DA neurons have been associated with decision-making under uncertainty ^18,30^, we first analyzed the spontaneous tonic activity of VTA DA cells in anesthetized mice. We recorded from mice chronically exposed to either saline or nicotine via minipump, and that had performed the behavioral task (“ICSS”, at the end of PS), or were behaviorally naïve. DA neuron firing was analyzed with respect to the average firing frequency and the percentage of spikes within bursts. As previously reported ^33,34^, chronic exposure to nicotine increased the tonic activity of DA neurons, both in terms of firing frequency and bursting activity, when compared to mice implanted with a saline minipump (Figure 2B). Furthermore, mice exposed to the ICSS task exhibited an increase in firing frequency, but no change in bursting activity when compared to mice that were not stimulated (Figure 2B).

**Figure 2:**
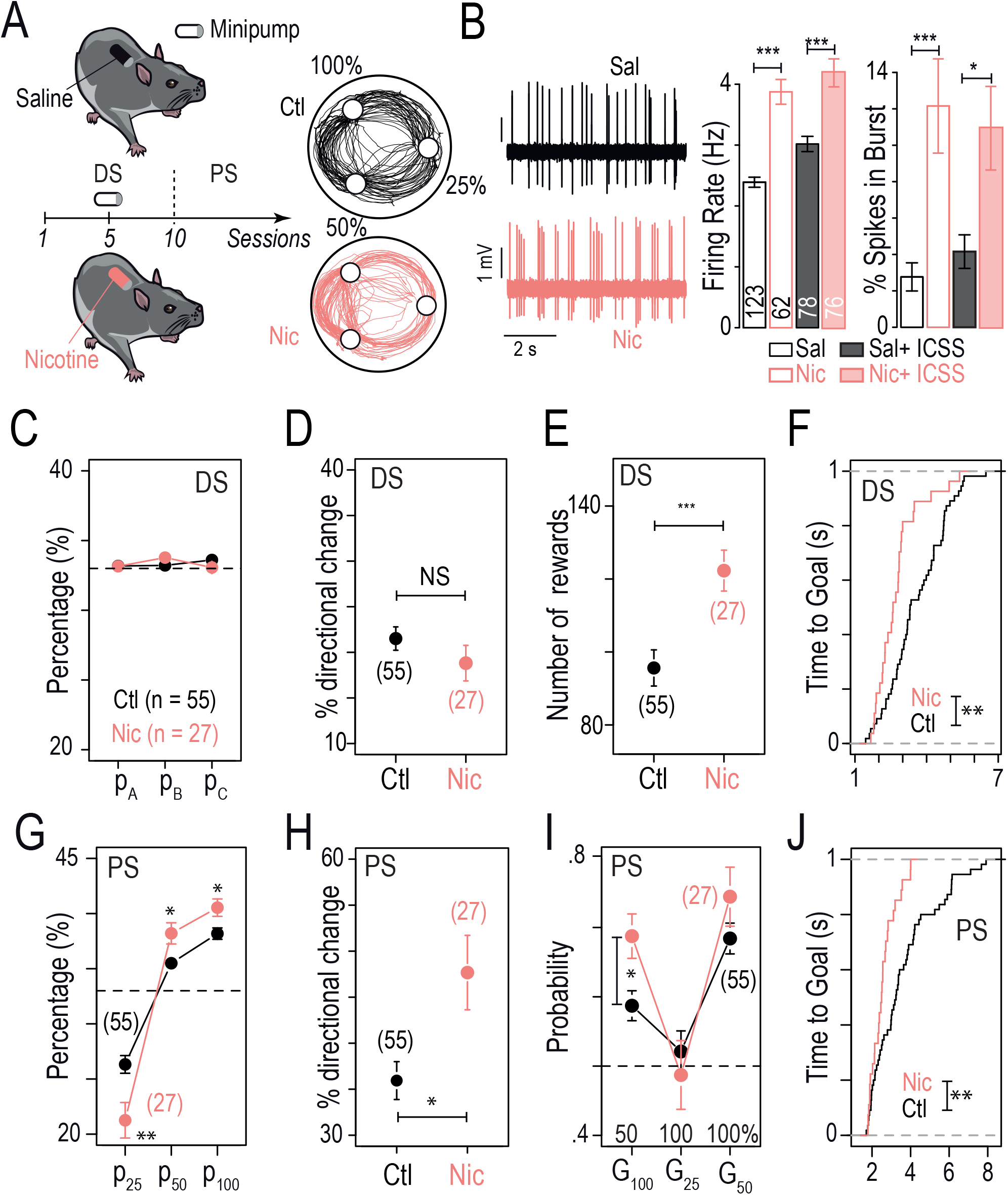
Chronic exposure to nicotine altered both spontaneous DA activity and choice strategies. (A) *Left*: Timeline of the task. Subcutaneous osmotic mini-pumps delivering nicotine (10 mg/kg/day), or saline for control animals, were implanted on day 5 of the DS. *Right*: Sample trajectories at the end of the PS for a mouse under chronic nicotine (Nic, in pink) and a mouse naive to nicotine (Ctl, in black). (B) *Left*: Representative electrophysiological recordings of VTA DA neurons after chronic saline (Sal, black) or nicotine (Nic, red) exposure. *Right*: The firing frequency and bursting activity of VTA DA neurons were compared between two sets of conditions: saline (n = 123) versus nicotine minipump (n = 62), and saline minipump + ICSS (n = 78) versus nicotine minipump + ICSS (n = 76) after completion of the PS. All electrophysiological experiments were performed after 24 ± 2 days of Sal or Nic (10 mg/kg/day) exposure. Nicotine exposure increased both DA neuron firing frequency (two-way ANOVA, nicotine effect *F*_(1,335)_ = 72.42, *p* < 0.001) and bursting activity (*F*_(1,335)_ = 25.39, *p* < 0.001), with or without ICSS. This increase was observed between the Sal and Nic minipump-only conditions (*post hoc* Tukey HSD, firing frequency p < 0.001, bursting activity p < 0.001), as well as after Nic minipump + ICSS compared to Sal minipump + ICSS (*post hoc* Tukey HSD, firing frequency p < 0.001, bursting activity p = 0.02). Mean firing frequency was increased after ICSS in both the Sal and Nic groups (two-way ANOVA, ICSS effect *F*_(1,335)_ = 21.53, *p* < 0.001), but bursting activity was unchanged after ICSS (*F*_(1,335)_ = 1.02, *p*= 0.31). No interaction effect was observed for firing frequency (*F*_(1,335)_ = 1.02, *p* = 0.31) nor bursting activity (*F*_(1,335)_ = 0.65, *p* = 0.42). (C-F) Comparison between mice exposed to chronic nicotine (Nic, in pink, n = 27) and control mice (Ctl, n = 55) at the end of the DS, regarding (C) the target repartition (i.e., P_A_, P_B_ and P_C_, p>0.05), (D) the percentage of directional changes (Student’s t-test, p>0.05), (E) the number of rewards (Student’s t-test, ***p < 0.001) and (F) the cumulative distribution of the average time-to-goal (KS test, **p < 0.01). (G-J) Comparison between mice exposed to chronic nicotine (Nic, in pink, n = 27) and control mice (Ctl, n = 55) at the end of the PS, regarding: (G) the target repartition. Nic mice visited more often the options with a higher reward probability (i.e. P_50_ and P_100_) and less often the option with the lowest probability (P_25_) in comparison to Ctl mice (student t-test with Holm correction for multiple comparisons, **p = 0.006, *p = 0.011, *p = 0.012, respectively). (H) Percentage of directional changes (Student’s t-test, *p = 0.02); (I) Probability of making the exploitative choice (i.e., the one with the highest probability of reward) for the three possible gambles for Nic and Ctl mice (Student’s t-test with Holm correction for multiple comparisons, *p = 0.03) and (J) the cumulative distribution of the average time-to-goal (KS test, **p < 0.01).

We then analyzed the behavior of mice in the ICSS task. Overall, we did not see any behavioral difference between mice implanted with a saline minipump (n=23) and the non-implanted mice (n=32) analyzed in Figure 1 (Supplementary Figure 2). Therefore, these two groups were pooled and henceforth referred to as control (Ctl, n=55). Trajectories at the end of the DS were stereotyped, almost circular, in both Ctl and Nic mice. Both groups distributed their visits equally over the three locations (Figure 2C) and their respective probabilities of directional changes were equal (Δ = −2.7% %, Figure 2D). However, the total number of rewards was higher for Nic mice than for Ctl mice (Δ = 26, Figure 2E), as a consequence of the decrease in the mean time-to-goal (i.e., the time necessary to go from one target to the next) in Nic mice (Δ = 0.83 s, Figure 2F). When mice were placed in a classical open field (without ICSS), a greater velocity was observed in mice exposed to nicotine, yet only at the beginning of the session (Supplementary Figure 3). This result suggests that the increased speed observed in the ICSS task for nicotine-treated mice may arise from the combined effects of nicotine exposure and the stimulation rewards.

Clear differences in the behavior of nicotine- and saline-exposed mice were observed in the PS. Both groups distributed their choices depending on the probability to receive a reward, but with different strategies. Notably, while Ctl mice visited significantly p_25_, Nic mice focused on the two most rewarded options (i.e., p_50_ and p_100_, Figure 2G, Δ_25_ = −5%, Δ_50_ = 2.7 %, Δ_100_ = 2.3%). These modifications were associated with an increase in the percentage of directional changes (Δ = 11%, Figure 2H) and in the optimal choice in gamble G_100_ (Figure 2I, Δ = 10%) for Nic mice compared to Ctl mice. We also observed an increase in the total number of obtained rewards (Δ = 17.9, p = 0.002) and in the percentage of success (number of rewards divided by the number of trials, Δ = 2 %, p = 0.02) in Nic mice compared to Ctl mice. Finally, the comparison of mean time-to-goal between the two groups (Δ = −1.1 sec, Figure 2J) indicates again an increased velocity in Nic mice, as was already observed in the DS. This increase in speed in the PS is not associated with a decrease in the number of directional changes made by Nic mice, suggesting that animals did not enter an automatic circular mode, disengaged from actual choices, but instead remained in a deliberative process. Altogether, these results indicate chronic nicotine modifies the decision strategy of mice by biasing choices toward the most immediately valuable options, and thus reduces exploration.

In the PS, adopting a purely exploitative strategy to maximize the success rate would require solely the alternation of visits between p_100_ and p_50_. Both Ctl and Nic groups clearly deviated from this strategy of pure exploitation, although Nic mice were more exploitative on average. Yet population analyses (i.e. averaging over groups of animals) classically do not reflect the wide range of distinct behaviors and strategies that can be adopted by individuals. We therefore further analyzed our behavioral data, with the aim of revealing individual profiles and their adaptation under nicotine exposure.

### Archetype analysis suggests that mice exhibit inter-individual variability in choice strategies, with chronic nicotine fostering exploitative profiles

Visual inspection of individual trajectories revealed that in the PS, some mice retained a circular strategy (with either an ascending (p_25_ - p_50_ - p_100_) or descending (p_100_ - p_50_ - p_25_) order) while others had what we hereafter call a gain-optimizing (GO) strategy, alternating between targets associated with the highest reward probabilities (p_100_ and p_50_ - Figure 3A, lower left). By a gain-optimizing strategy, we mean a very basic definition of optimality based only on maximizing the number of rewards, but which does not take into account the advantage of exploration. Theoretically, always choosing the most valuable option would lead to an average success rate of 75% (Figure 3A, lower right) while a purely circular strategy would lead to an average estimate of 58.3% success rate (Figure 3A, upper right). Accordingly, the percentage of directional changes was correlated with the success rate (Figure 3B, for Ctl and Nic mice). Progressively adding directional changes between the p_50_ and p_100_ targets to the circular pattern theoretically result, in this plot, in a displacement along the line that connects the theoretical points of the circular strategy (0% U-turn, 58.3% success) to the gain-optimizing strategy (100% directional changes, 75% success) (Figure 3B, red line). We found experimentally that the slope (s = 17.1 ± 1.5, black line, Figure 3B) of the correlation between directional change and Success rate was almost parallel to the theoretical line from circular to gain-optimizing strategies (S_th_= 16.7, red line, Figure 3B), indicating that most of the directional changes were not random, but consisted in back-and-forth sequences between the p_50_ and p_100_ targets.

**Figure 3:**
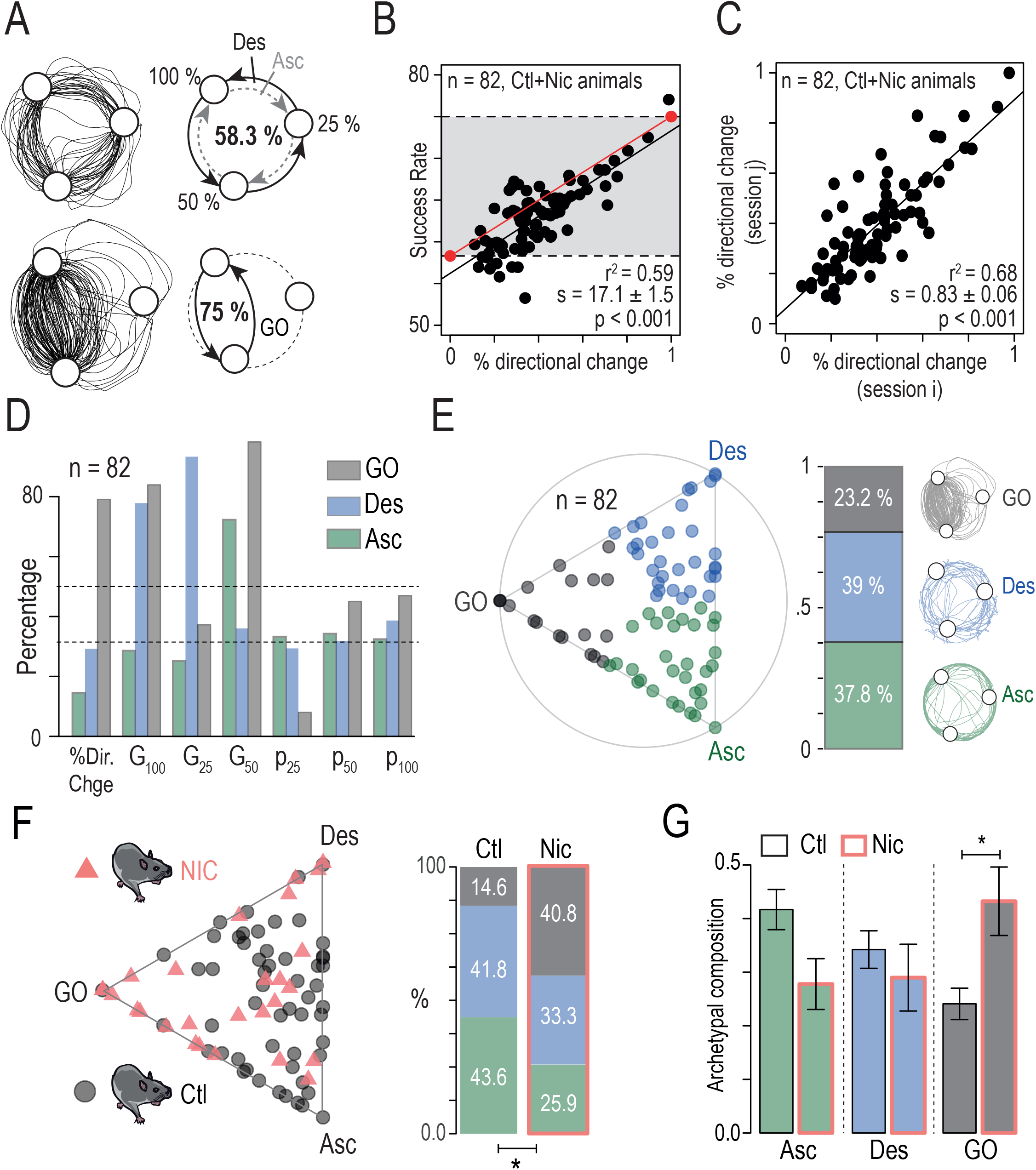
Mice exhibited inter-individual differences in choice strategies which were differentially affected by chronic nicotine exposure. (A) *Left*: Sample trajectories in the PS, corresponding to different choice strategies, a circular strategy (*top*) and a gain-optimizing strategy (*bottom*). *Right*: A mouse using a purely circular strategy (top, descending Des or ascending Asc) in the PS will tend to a 58.3 % success rate, whereas a mouse that always avoids p_25_ and alternates between p_100_ and p_50_ (bottom) will reach 75 % of success rate. (B) Correlation between the success rate and the percentage of directional changes. Mice displayed a strong inter-individual variability in their choice strategy but, overall, the higher the percentage of directional change, the higher the success rate (regression line in black). The red line indicates the linear correlation passing through two theoretical points: {0% directional changes; 58.3 % success rate} and {100 % directional changes; 75 % success rate}. (C) Correlation between the percentage of directional changes for two consecutive sessions. This measure showed a strong stability between consecutive sessions, indicating that the decision strategy was conserved across time for a given individual. (D-E) Archetypal analysis of the choice strategies based on 7-dimensional data space: i) the % of directional changes, ii) the gambles G_100_, G_25_ and G_50_, and iii) the distribution of choices between p_25_, p_50_, and p_100_. Analysis was performed on n = 82 mice (pooled Ctl and Nic mice). (D) Plot of the three archetypal solutions, gain-maximizers (GO), descending (Des) and ascending (Asc), and their 7 basic variables used in this analysis. (E) *Left*: Visualization of the α coefficients using a ternary plot. Each point represents the projection of an individual onto the plane defined by a triangle where the three apices represent the three archetypes (GO, Des, and Asc). Points are color-coded according to their proximity to the archetypes. *Right*: Proportions of each archetype on the entire population: 37.8 % Asc (green), 39 % Des (blue) and 23.2 % GO (grey). (F) *Left*: NIC (pink triangles) and Ctl (grey dots) mice displayed on the same ternary plot. Nic mice displayed a visual shift of their behavior towards the GO extrema of the archetype. *Right*: This shift was reflected by a difference in the proportion of each phenotype between Nic and Ctl groups (χ^2^ test, p = 0.027), with a higher proportion of GO mice in the Nic group. (G) Archetypal composition for each archetype (1 = closer to the apex) in Ctl and Nic mice (Wilcoxon test, p = 0.08, p = 0.22 and p = 0.04, with Holm correction).

To test whether the variabilities in behavior were robust for each individual from trial to trial, we compared the percentage of directional changes for two consecutive sessions for each animal of the Ctl group. Directional changes showed a strong positive correlation from one session to the next (Figure 3C), suggesting a strong consistency in individual behaviors and inter-individual variations in the strategy used during the PS. We thus characterized individual behaviors of all mice (both Ctl and Nic groups, i.e n=82) in the task using a seven-dimensional dataset based on the statistics of i) the directional changes, ii) the target distributions and iii) the three gambles (see data Figure 1 D-F). Principal-component analysis methods have been classically used to split high-dimensional data sets into clusters. Rather than aggregating individual data onto typical observations (the cluster centers), archetypal analysis ^35,36^ depicts individual behavior as a continuum within an “archetypal landscape” defined by extreme strategies, the archetypes. Individual data points are represented as linear combinations of extrema (vertex corresponding to “archetypal strategies”) of the dataset. The seven-dimensional dataset was used to identify three archetypal phenotypes. The three archetypes and their characteristics (Figure 3D) differentiated mice exhibiting a gain-optimizing strategy (i.e. focusing on p_50_ and p_100_), Figure 3A, below) which are referred to as gain-optimizers (GO, in grey), from mice with circular patterns (equal distribution between the three targets, Figure 3A above), which either turned in a descending manner (labelled Des, in blue, sequence p_100_ - p_50_ - p_25_ associated with high G_100_ and G_25_ but low G_50_) or an ascending manner (labeled Asc, sequence p_25_ - p_50_ - p_100_ associated with low G_100_ and G_25_ but high G_50_). The individual behavior of each of the 82 mice could be defined as a weighted combination of these three extremes in a ternary plot (Figure 3E). An animal’s behavior in this ternary plot is defined by three coordinates (a,b,c) that sum to 1 and that depict its relative archetypal composition. Therefore, these coefficients (a,b,c) could be used to assign an individual to its nearest archetype based on its behavioral profile (Figure 3E left). This assignment revealed that 23.2 % of the mice were closer to the GO archetype (grey), while the remaining mice were evenly distributed between the Des (39%, blue) and Asc archetypes (37.8%, green) (Figure 3E Right). To analyze the effect of chronic nicotine, we split Ctl and Nic mice, and showed that these two groups distributed differently in the archetypal space as indicated by a modification of i) the distribution of the archetype’s assignments (Figure 3F) and of ii) the archetypal composition (Figure 3G). Overall, chronic nicotine exposure produced an apparent displacement of the population further from Asc and Des apices and closer to the GO apex, thus it favored the emergence of the more exploitative, and thus less explorative, GO phenotype.

### Nicotine exposure modified decision parameters associated with exploration and cost

To quantitatively describe the effects of nicotine on the decision processes underlying choice behavior in mice, we modeled our data using a softmax model of decision-making. In this model, the probability of choosing a target A over B depends on the difference between their expected values, here the probability *p* of reward delivery associated with each target (as the stimulation magnitudes were the same for all targets), and the “inverse temperature” parameter β which represents the sensitivity to the difference of values (ΔV). A small β favors exploration (the proportion of respective choices is less sensitive to ΔV, with a null β meaning all options have nearly the same probability to be selected, independently of their respective value), while a large β indicates exploitation (high sensitivity to ΔV, with an infinite β meaning that options associated with higher reward probabilities are always selected). β can thus be considered as a proxy to measure the exploration/exploitation tradeoff. This model was adapted to account for the behavior of mice in the PS as follows: first, decisions were biased towards actions with the most uncertain consequences, by assigning a bonus value φ to the expected uncertainties, i.e. the variance *p*(1-*p*) associated with each location ^30^. This allowed us to explain the atypically low probability of choosing p_100_ over p_50_ in G_25_ (Figure 1F). Second, to account for the circular bias observed in both DS and PS, we added a motor cost which decreases the value of a target if it requires the animal to perform a directional change ^37^. Thus, in this adapted softmax model (Figure 4A and Methods), the “exploration/exploitation” parameter β represents how the probability to choose between options depends on the difference of their respective subjective values, which was defined as the weighted sum of the expected values (100, 50 or 25 %), expected uncertainty (weighted by parameter φ) and expected motor cost (weighted by parameter κ) of a given target.

**Figure 4:**
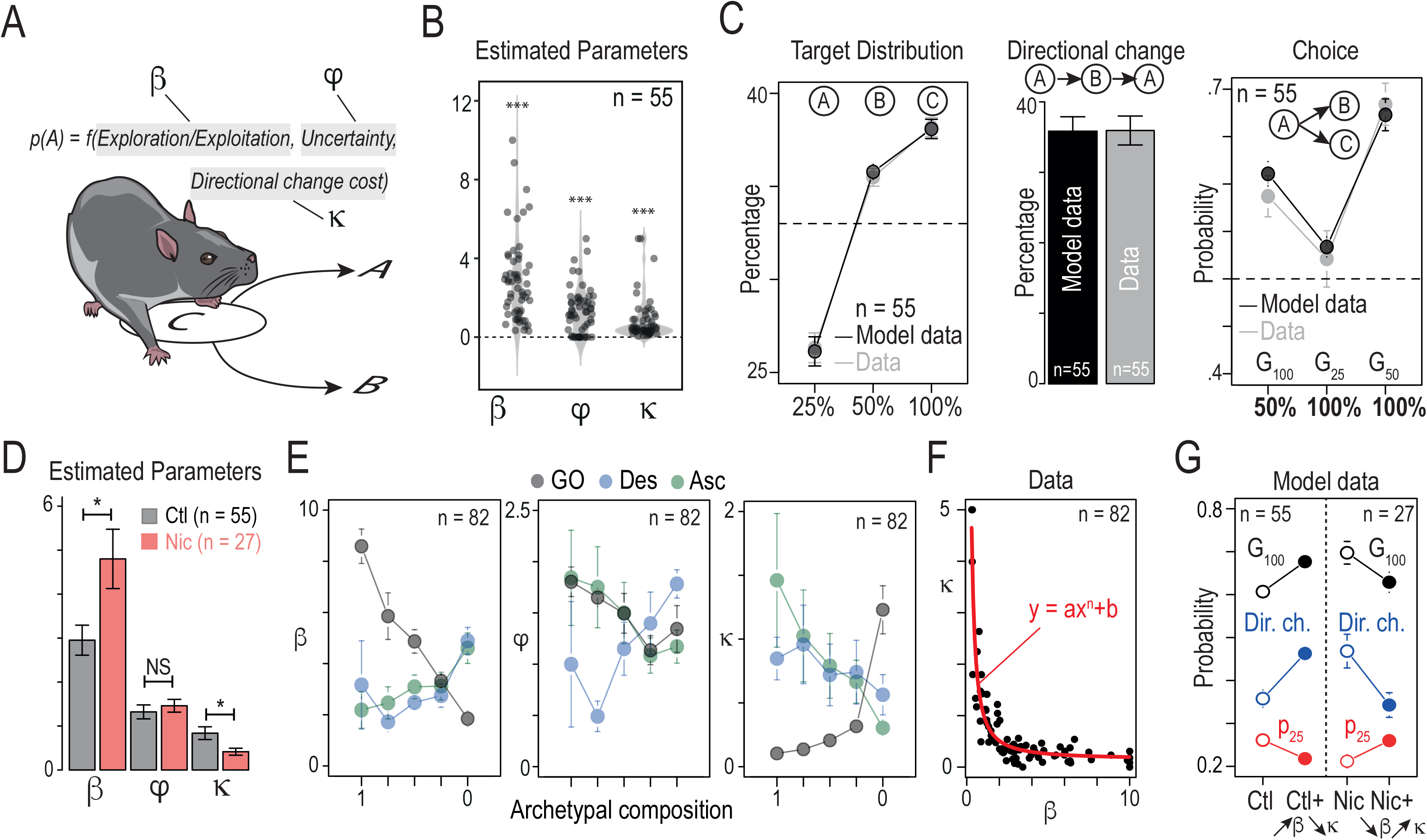
Computational modeling suggests that decision parameters differ between the three archetypes and are differentially affected by nicotine exposure. (A) Principle of the softmax model: softmax decision rule with three parameters β (inverse temperature or exploration/exploitation), φ (uncertainty bonus) and κ (cost or effort for a directional change). (B) Estimated values of β, φ and κ parameters for the 55 Ctl mice (not exposed to nicotine, *** indicates a significant difference from zero). (C) Comparison between Ctl data and model for a model sequence of 2000 choices (n = 55) simulated with fitted values of β, φ and κ (see B) *Left*: Repartition of visits on the three targets (p_25_, p_50_ and p_100_, with a mean of the differences between Ctl data and model of Δ = 0.002%, −0.003% and 0.001%, p > 0.05). *Middle*: Comparison of the percentage of directional changes (Δ = 0.002, p > 0.05). *Right*: Probability to choose alternatives with the highest probability of reward for the three possible gambles (G_100_ = p_50_ over p_25_; G_25_ = p_100_ over p_50_; G_50_ = p_100_ over p_25_, Δ = −0.02, −0.008 and 0.009, p > 0.05 for the three gambles). (D) Nicotine-exposed animals displayed an increase in β (Δ = 1.85, p = 0.03), a decrease in κ (Δ = −0.42, p = 0.04), but no difference in φ (Δ = 0.14, p > 0.05) compared to Ctl mice (Student’s t-test with Holm correction for multiple comparisons). (E) *Left:* Correlation between β (left), φ (middle) or κ (right) values and the archetypal composition for both Ctl and Nic mice (n = 82, see plot Figure 3B). The closer to the GO phenotype, the higher the β and the lower the κ, which is consistent with an optimal strategy based on alternation between p_100_ and p_50_. The closer to the Des phenotype, the lower the φ parameter. (F) Plot of the fitted β and κ parameters for both Ctl and Nic mice (n = 82). Data are fitted with a polynomial function (y = ax^n^+b) (G) Mimicking the effect of nicotine on the model parameters. *Left*: The simulation of choice behavior when nicotine-induced increase of β and decrease of κ are added to the Ctl model parameters (n = 55, Ctl + ↗β↘κ) recapitulates the effect of nicotine on the three choice parameters (the probability to choose the most valuable option in gamble G_100_; the percentage of directional changes, and the probability to visit p_25_, mean of the differences between Nic data and model Δ = 0.01%, −0.005%, 0.01%, respectively, Student’s t-test, p > 0.05). Starting from the Nic mice parameters and removing the nicotine-induced changes on β and κ (n = 27, Nic + ↘β↗κ) reestablish those three parameters at the level of Ctl mice (mean of the differences between Ctl data and model Δ = 0.02%, −0.0006%, −0.03%, respectively, Student’s t-test, p>0.05). Δβ is calculated using β_Nic_ – β_Ctl_ the mean estimated in Ctl and Nic condition. κ is determined using the non-linear relationship between β and κ (see F).

We fitted the transition function of each mouse from the Ctl group (n = 55) with this model, resulting in positive β, φ and κ values (Figure 4B). The robustness of the model was then assessed by generating sequences of choices (n = 2000 model choices) for n = 55 mice with their respective model parameters (Figure 4C). The model accurately reproduced the mean distribution of targets (Figure 4C, Left), the proportion of directional changes (Figure 4C, Middle) and the choice transition function (Figure 4C, Right). Individual transition functions from Nic mice (n=27) were then fitted by the same model. When compared with the model parameters of Ctl mice, nicotine exposure increased the value sensitivity parameter β, and decreased the cost of directional changes κ parameter, but did not affect the uncertainty bonus φ (Figure 4D).

We then assessed whether archetypal phenotypes of choice data could be derived from differences in decision-making processes measured by the parameters of our model by evaluating the value of the three parameters (β, φ, κ) depending on the archetypal composition (see methods). Overall, the three archetypes corresponded to different combinations of the model parameters (Figure 4E). The GO (grey) archetype was associated with a high value of β (corresponding to exploitation) and φ but a low motor cost κ, which is consistent with individuals that favor the alternation between locations associated with higher probability (p_100_ and p_50_). The Des and Asc phenotypes corresponded to strong circular behaviors and thus to high motor cost κ and low sensitivity to value β. Des and Asc differed by their φ value (Δ = 1.012, p = 0.0079), which was related with the directionality of their preferred rotation: a low preference to uncertainty φ corresponds to mice choosing the certain p_100_ reward over the uncertain p_50_ reward, resulting in a tendency for sequence p_25_ -> p_100_ -> p_50_ observed in Des mice (blue). Conversely, a high preference for uncertainty φ is associated with the reverse sequence p_25_ -> p_50_ -> p_100_ observed in Asc mice (green). β and κ appeared non-linearly correlated, as indicated by the negative relationship between archetypal composition and β and κ variations (Figure 4E) and by the inverse correlation between β and κ (Figure 4F, pooled Ctl and Nic groups). Overall, the decomposition of the archetypal phenotypes into their underlying decision-making processes illustrate how distribution of individual decision-making strategies (Asc, Des and GO) in a population, could corresponds to transitions in the parameter values from the same model. It also allows us to interpret the effects of nicotine as a coordinated increase of β and decrease of κ consistent with a deviation towards the GO profile. We thus asked whether recapitulating these effects on decision parameters β and κ would be sufficient to shift decision-making strategy towards the GO profile. To test this idea, we modeled the choices (N = 2000) using decision-making parameters from the Ctl population (n=55, as in Figure 3B-C) modified by the average difference observed in the β and κ parameters from Nic mice. To avoid spurious effects associated with the resulting β and κ values falling outside the range observed for Nic + Ctl mice, we increased the β parameter and derived κ from their non-linear relationship (fitted in Figure 4F). We evaluated the consequences of mimicking the effect of nicotine on decision-making parameters by comparing the three main behavioral measures altered by nicotine: i) the probability to choose the most valuable option in gamble G_100_ (choosing p_50_ over p_25_), ii) the percentage of directional changes and iii) the probability to visit p_25_. By applying a combination of β increase and κ decrease (derived from Nic mice) to the Ctl model parameters, the model accurately reproduced, for the three measures (Figure 4G), the changes observed in decision-making strategy following chronic nicotine exposure. Conversely, by combining a decrease in β with an increase in κ (i.e. subtracting the average effect of nicotine from the Nic model parameters) we are able to simulate the conversion of a Nic behavioral profile into a Ctl profile.

### Optogenetic stimulation of VTA DA neurons recapitulated the effects of nicotine

Finally, to assess whether the changes we observed in decision-making strategies following chronic nicotine exposure could be linked to the alterations of VTA DA neuron activity, we sought to experimentally alter choice behaviors by acutely manipulating the activity of VTA DA neurons using optogenetics. Nicotine exposure is known to induce modifications in a number of brain areas ^38^, including an increase in the tonic activity of VTA DA neurons, as we indeed found in this study (Figure 2B). Furthermore, the tonic activity of DA neurons has been proposed to play a role in the balance between exploration and exploitation ^26–28^. We thus asked whether directly modifying the firing pattern of VTA DA neurons was sufficient to alter decision-making behavior and recapitulate the effects of chronic nicotine in our ICSS task. To specifically and bi-directionally manipulate VTA DA neurons, we expressed either an excitatory channelrhodopsin (CatCh, ^39^) or an inhibitory halorhodopsin variant (Jaws, ^40^) in DAT^iCRE^ mice using a Cre-dependent viral strategy (Supplementary Figure 5A). We confirmed in patch-clamp recordings that continuous 5ms-light pulses at 8 Hz (470 nm) reliably increased VTA DA neuron activity in CatCh-transduced mice (Figure 5A), while 500ms-light pulses at 0.5 Hz (520 nm) reliably decreased their activity in Jaws-transduced mice (Figure 5B).

**Figure 5:**
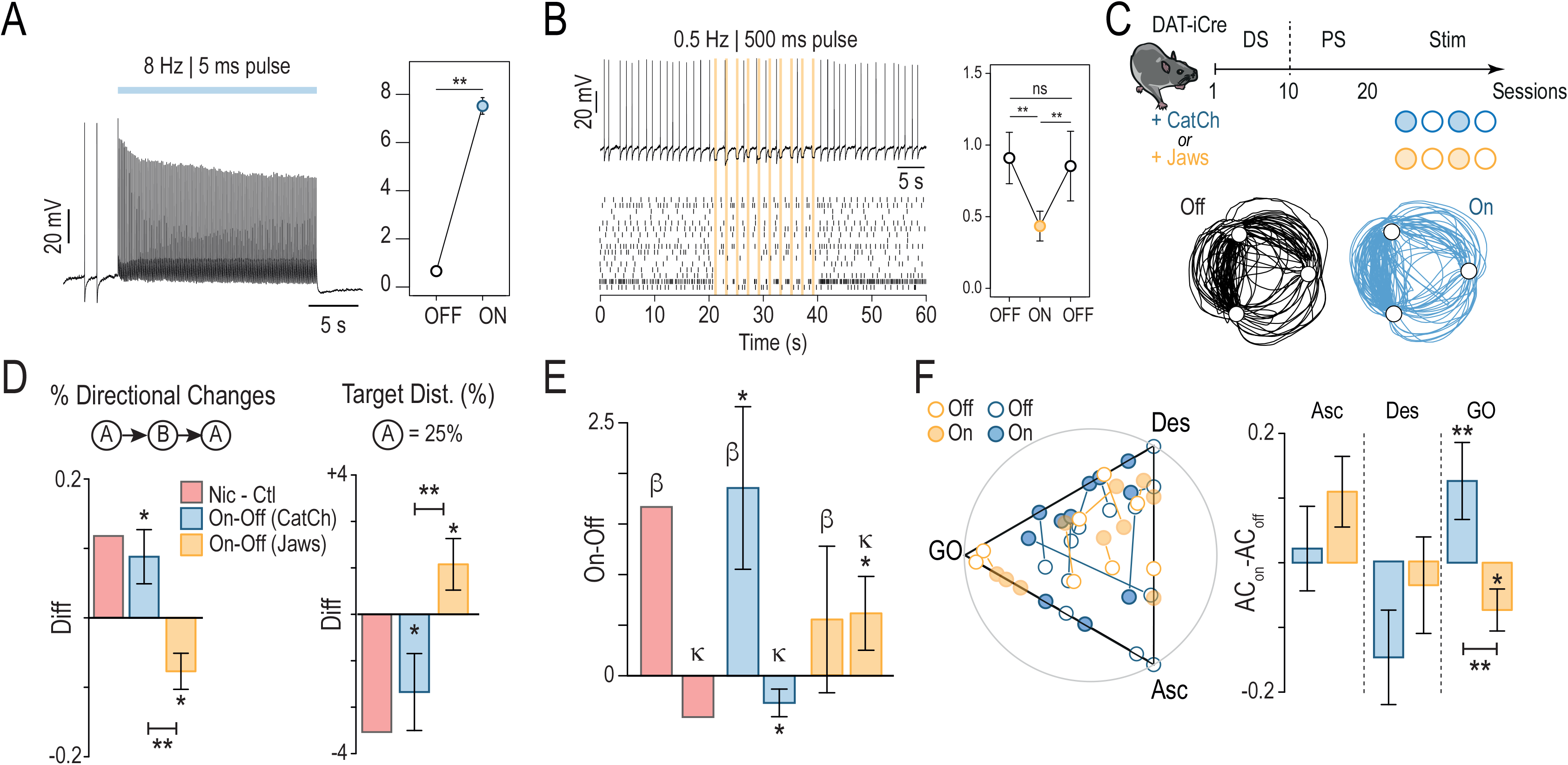
Optogenetic manipulation of VTA DA neuron activity recapitulated the behavioral adaptations observed under chronic nicotine exposure. (A) *Left:* representative current-clamp recording of a VTA DA neuron transduced with CatCh and stimulated with 5-ms blue light pulses at 8 Hz. *Right*: Average increase in basal firing frequency upon optogenetic stimulation for n = 10 neurons (p-value = 0.002, Wilcoxon test). (B) *Top left:* representative current-clamp recording of a VTA DA neuron transduced with Jaws and stimulated with 500-ms green light pulses at 0.5 Hz. *Bottom left:* Raster plot for n = 16 neurons. *Right*: Average decrease in basal firing frequency upon optogenetic stimulation, and return to the baseline after the photostimulation period, for n = 16 neurons (p-value: ns = 0.18; **0.004; **0.0014, Wilcoxon test with Holm correction). (C) Task design and photo-stimulation protocols. DAT-iCre mice transduced with either an AAV-DIO-Catch-YFP in the VTA (CatCh, blue) or an AAV-DIO-Jaws-eGFP (Jaws, yellow) and were implanted unilaterally with bipolar stimulating electrodes for ICSS in the MFB. Following the DS and PS sessions they received 2 paired ON (*filled circles*) and OFF (*open circles*) sessions with the same rules as PS. *Below:* Representative trajectories of a CatCh-transduced mouse with (*blue*) and without (*black*) optogenetic stimulation of VTA DA neurons. (D) Net effect of light stimulation for the percentage of directional changes (*left*) and the proportion of p_25_ visits (*right*). Data from OFF sessions were subtracted from data from ON sessions. In red, the net effect of nicotine was represented for all the parameters, as a comparison factor. (Asterisk: Comparison with a true mean of 0 and paired comparison (On-Off), Student’s t-test or Wilcoxon test, unilateral testing). (E) Net effect of photo-stimulation on the softmax model parameters β and κ (Student’s t-test or Wilcoxon test, unilateral testing). (F) *Left*: Position of each animals in the ternary archetype plot. *Right*: difference in archetypal composition (ON-OFF) for each archetype. Optogenetic activation of DA neurons triggered a shift of the behavior towards the GO phenotype while optogenetic inhibition induced a shift of the behavior away from GO.

After mice completed both the DS and PS in the ICSS task, they went through four stimulation sessions (Stim) maintaining the same rules as the PS, with an alternating schedule of two days with photo-stimulation (ON, photo-stimulation started 5 min prior to the start of task and was maintained throughout) and two without (OFF) (Figure 5C). During the OFF days, mice were connected to the optical fiber patch-cord but did not receive light stimulation. For each pair of ON/OFF experiments, we estimated the effect of the photo-stimulation on the four main behavioral measures that were altered by chronic nicotine by calculating for a given measure (M) the difference M_ON_-M_OFF_ (Figure 5D, Supplementary Figure 5B-C). Overall, we found that optogenetic activation and inhibition of VTA DA neurons had opposite effects on behavioral outcomes such as the time to goal (Supplementary Figure 5B), the proportion of directional changes (Figure 5D, left), the proportion of visits on p_25_ (Figure 5D right), and the choice made in gamble G_100_ (Supplementary Figure 5C). Optogenetic activation increased directional changes (Figure 5D left) and decreased the probability to visit p_25_ (Figure 5D right), favoring alternations between p_100_ and p_50_, similar to the effect of nicotine. Opposite effects were observed for these two parameters when the firing rate was reduced in VTA DA cells using Jaws (Figure 5D). However, such optogenetic inhibition did not significantly affect the time to goal (Supplementary Figure 5B) nor the choice in the gamble G_100_ (Supplementary Figure 5C).

We then fitted the transition function of Catch- and Jaws-transduced mice with our decision-making model. As we observed with chronic nicotine, the effects of photo-activating VTA DA neurons on decision-making within the ICSS task could be modeled as an increase of β and a decrease of κ (Figure 5E). Photo-inhibition of VTA DA neurons, however, produced an apparent increase in the motor cost κ, opposite to the effect of chronic nicotine, but with no significant effects on the exploration/exploitation tradeoff parameter β (Figure 5E). By analyzing decision-making behaviors between the stimulated (ON) and non-stimulated (OFF) conditions in the previously identified archetypal space, we revealed that VTA DA neuron activation draws individual phenotypes towards the GO archetype (i.e, increased GO archetypal composition), while VTA DA neuron inhibition drew individuals away from GO (Figure 5F). Thus, altering the firing pattern of VTA DA neurons, by changing both the motor cost and the balance between exploration and exploitation behavior, is sufficient to drive the bias of decision behaviors in the ICSS task, as suggested by our simulations (Figure 4). Furthermore, increasing VTA DA neuron firing mimicked the effects of chronic nicotine exposure on decision-making measures, linking the behavioral alterations with the physiological changes to DA neurons we observed in Nic mice.

## Discussion

Understanding how nicotine affects decision-making has been challenging, because two different physiological aspects need to be distinguished ^31^: (1) nicotine as a reinforcer that directly activates the dopaminergic system to produce reinforcement and nicotine-seeking, and (2) nicotine as a neuromodulator that alters nicotine-independent decision-making processes by modifying the dynamics and computational properties of cholinoceptive circuits. Here, using a multi-armed ICSS bandit task, we show that mice passively treated with nicotine progressively learn to forage more frequently at locations with the highest probabilities of reward (p_50_ and p_100_) compared to naive animals, suggesting a bias in the exploration/exploitation tradeoff which decreases exploration. Acutely increasing the tonic activity of VTA DA neurons during the task recapitulated the effects of chronic nicotine exposure on mouse decision-making.

In our experiment, mice adapted their choices according to the probability of reward delivery, but they also consistently continued to visit the targets associated with lower reward probabilities in all of the gambles, even after extended training. Such a high level of exploratory behavior is potentially attributable to the setup, which is characterized by the delivery of small rewards, serially repeated gambles with short delays between trials, and learning through experience ^41^. The fact that mice continue to visit targets with the lowest probability in each of the gambles, despite intensive learning, can reflect i) exploratory noise, generally modeled via decreased value sensitivity (or increased randomness) of β in the softmax model, ii) directed exploration, if one considers that mice continue to explore locations associated with low reward probability to reduce the uncertainty associated with probabilistic omission, and iii) uncertainty-seeking, which is neither explorative nor exploitive but considers that mice simply attribute a positive value to expected uncertainty. Mouse choices and qualitative inter-individual variations were well described by a simple computational model of decision-making that takes into account exploration/exploitation tradeoff, uncertainty, and motor cost. Idiosyncrasy in choice behavior was well reflected by continuous variations in the key parameters of this model. Despite variations in individual choice behaviors, the consequences of nicotine administration were consistent, with a clear effect on the β and κ parameters, and a strategy biased towards the exploitation of the highest reward values.

The increase of β reflects an amplified exploitative behavior, an effect that has been previously linked to enhanced tonic DA activity, which is hypothesized to modulate the bias towards optimal choices ^26–28^. In this study, we demonstrate a direct link between DA neuron tonic activity and exploitation using electrophysiological and optogenetic approaches. The multi-armed ICSS bandit task enables, through a clear distinction between action selection (choices) and action execution (time to goal), to identify the modified components of value-based decision-making in relation to tonic DA. We explicitly demonstrate an increase in value sensitivity due to nicotine-induced alterations in tonic DA activity. Previous ICSS studies have observed that chronic exposure to drugs sensitizes the brain reward system, and in doing so lowers the stimulation threshold (expressed as a current intensity or stimulation frequency) ^42^ required for ICSS ^43^. Here we expand these results by quantifying the effects of such increased value sensitivity on choices between ICSS-mediated rewarding locations, and further identifying a causal link between these behavioral modifications and increased tonic activity of VTA DA neurons. Long-term nicotine exposure increases the basal activity of VTA DA neurons ^33,34^ through desensitization and up-regulation of nAChRs and the long-term strengthening of glutamatergic synaptic transmission ^44^. Here we show that elevating VTA DA neuron activity in an acute fashion using optogenetics is sufficient to induce behavioral alterations in mice similar to those that we observed following chronic nicotine exposure.

Variations in neuromodulatory functions, including those in the catecholamine and cholinergic systems, contribute to the process of individuation ^45–47^. DA, and in particular from VTA DA neurons, has been linked to a cluster of traits (extraversion, novelty-seeking, etc.) conceptually related to reward-seeking ^48 49^. However, despite the substantial attention paid to DA in personality neuroscience, and despite a clear association between modulations in dopaminergic function and variations in individual traits, defining which specific traits are influenced by DA remains a challenging task. Our data suggest that modification in basal VTA DA neuron activity can directly modify the expression of one central personality trait: exploration. This result is reminiscent of the observations made from mice living continuously in a large environment, which display idiosyncratic behavioral strategies during a decision-making task, and for which the exploration/exploitation balance was correlated with the activity of their DA system^47^.

Nicotine exposure alters decision-making processes ^6^. Non-contingency studies have previously shown that yoked nicotine exposure increases the incentive salience of non-nicotine stimuli ^50^, similar to the sensitization to ICSS rewards ^43^. These studies suggest an essential role of contextual cues in smoking and the nicotine-induced increase in reward sensitivity. Neuroeconomics studies have also linked smoking with increased impulsivity (delay discounting task ^8^), lack of counterfactual learning signals ^51^, and decreased behavioral flexibility (exploration in a dynamic bandit task ^10^). Our results further reveal that nicotine exposure decreases exploration. In addition, we provide a mechanistic understanding of how reward processing may be altered at the level of the VTA in smokers. Our data underscore altered choice behaviors in smokers that likely participate in, but are not limited to, addiction^6^. Nicotine-induced alterations in decision-making processes likely also have implications for everyday life, particularly as they can increase vulnerability for addiction to other drugs of abuse and for behavioral disorders such as pathological gambling that rely on value-based decisions ^7,52^ and present a high comorbidity with tobacco addiction ^53^.

## Acknowledgements

We are grateful to the animal facilities (IBPS), Camille Robert and Paris Vision Institute AAV production facility for viral production and purification. This work was supported by the Centre National de la Recherche Scientifique CNRS UMR 8246, INSERM U1130, the Foundation for Medical Research (FRM, Equipe FRM DEQ2013326488 to P.F), FRM FDT201904008060 (to SM), the French National Cancer Institute Grant TABAC-16-022 et TABAC-19-020 (to P.F.), French state funds managed by the ANR (ANR-16 Nicostress to PF, ANR-19 Vampire to FM) and The LabEx Bio-Psy (to P.F). MLD, RDC and SM were the recipients of a fourth-year PhD fellowship from FRM (FDT20160435171, FDT20170437427 and FDT201904008060), CN was recipient of a doctoral fellowship from the Labex Bio-Psy, DL was recipient of a post-doctoral Fellowship from the Labex Bio-Psy, and LMR was supported by a NIDA–Inserm Postdoctoral Drug Abuse Research Fellowship.”

## Author contributions

PF and MD designed the study. MD, RDC, CN, MC, TAY, EKD, RB, EB, BH, and NT performed the behavioral experiments. MD, RDC, SM and NT performed the minipumps implantations. JN, DL and SD contributed to setup developments. CN, SM, RDC, DL and FM performed electrophysiological recordings. MD, RDC, CN, MC, EKD, RB, TAY, EB, NT and JN performed the surgeries and virus injections. CN, SM, performed the immunohistochemistry experiments. DD provided the viruses. JN and PF developed the model. AM developed the optogenetic setup. MD, RDC, CN, MC, SM, JN, FM and PF analyzed the data. PF wrote the manuscript with inputs from MD, RDC, CN, MC, LMR, JN, FM and AM.

## Methods

### Animals

Experiments were performed on adult C57Bl/6Rj DAT^iCRE^ and Wild-Type (Janvier Labs, France) mice. Male mice, from 8 to 16 weeks old, weighing 25-35 grams, were used for all the experiments. They were kept in an animal facility where temperature (20 ± 2°C) and humidity were automatically monitored and a circadian light cycle of 12/12-h light-dark cycle was maintained. All experiments were performed in accordance with the recommendations for animal experiments issued by the European Commission directives 219/1990, 220/1990 and 2010/63, and approved by Sorbonne University.

### AAV production

AAV vectors were produced as previously described using the cotransfection method and purified by iodixanol gradient ultracentrifugation^51^. AAV vector stocks were tittered by quantitative PCR (qPCR)^52^ using SYBR Green (Thermo Fischer Scientific).

### Intracranial self-stimulation electrode implantation

Mice were anaesthetized with a gas mixture of oxygen (1 L/min) and 1-3 % of isoflurane (Piramal Healthcare, UK), then placed into a stereotaxic frame (Kopf Instruments, CA, USA). After the administration of a local anesthetic (Lurocain, 0.1 mL at 0.67 mg/kg), a median incision revealed the skull which was drilled at the level of the Median Forebrain Bundle (MFB). A bipolar stimulating electrode for ICSS was then implanted unilaterally (randomized) in the brain (stereotaxic coordinates from bregma according to mouse after Paxinos atlas: AP −1.4 mm, ML ±1.2 mm, DV −4.8 mm from the brain). Dental cement (SuperBond, Sun Medical) was used to fix the implant to the skull. After stitching and administration of a dermal antiseptic, mice were then placed back in their home-cage and had, at least, 5 days to recover from surgery. An analgesic, buprenorphine solution at 0,015 mg/L (0,1 mL / 10 g), was delivered after the surgery and if necessary, the following recovering days. The efficacy of electrical stimulation was verified through the rate of acquisition during the deterministic setting (see behavioral methods).

### Implantation of osmotic mini pumps

After 5 days of training in the deterministic setting (see behavioral methods), animals were anesthetized with a gas mixture of oxygen (1L/min) and 1-3 % of isoflurane (IsoVet, Piramal Healthcare, UK). After the administration of a local anesthetic, an incision was performed at the level of the interscapular zone, to subcutaneously implant an osmotic minipump (Model 2004, ALZET, CA, USA) containing 200 μL of either a solution of nicotine hydrogen tartrate salt (Sigma-Aldrich, USA) at a dose of 10 mg/kg/d or saline solution (H_2_O with 0.9 % NaCl) for the control group. Both solutions were prepared in the laboratory. Minipumps delivered their content with a flow rate of 0.25 μL/hour over 28 days. The surgical wound was closed with surgical stitches. Animals had two days of rest to recover from the minipump surgery before going further with their behavioral training.

### Virus injections and optogenetics experiments

DAT^iCRE^ mice were anaesthetized (Isoflurane 1-3%) and implanted with an ICSS electrode as described above. They were then injected unilaterally (randomized left/right side and ipsi/contralateral side regarding the ICSS electrode) in the VTA (1 μL, coordinates from bregma: AP −3.1 mm; ML ±0.5 mm; DV −4.55 mm from the skull) with an adeno-associated virus (AAV5.EF1α.DIO.hCatCh.YFP 2.46e^12^ - 6.53e^13^ ng/μL, AAV5.EF1α.DIO.Jaws.eGFP 1.16e^13^ ng/μL or AAV5.EF1α.DIO.YFP 6.89e^13^ or 9.10e^13^ ng/μL). A double-floxed inverse open reading frame (DIO) allowed to restrain the expression of CatCh (Ca^2+^-translocating channelrhodopsin) or Jaws (red-shifted cruxhalorhodopsin) to VTA dopaminergic neurons.

For optogenetic experiments on freely moving mice, an optical fiber (200 μm core, NA = 0.39, Thor Labs) coupled to a ferule (1.25 mm) was implanted just above the VTA ipsilateral to the viral injection (coordinates from bregma: AP −3.1 mm, ML ±0.5 mm, DV 4.4 mm), and fixed to the skull with dental cement (SuperBond, Sun Medical). The behavioral task began at least 4 weeks after virus injection to allow the transgene to be expressed in the target dopamine cells. An ultra-high-power LED (470 nm or 520 nm, Prizmatix) coupled to a patch cord (500 μm core, NA = 0.5, Prizmatix) was used for optical stimulation (output intensity of 10 mW). Optical stimulation was delivered continuously, starting 5 min before and continuing throughout the 5 min of ON sessions of the task. Excitatory opsin (CatCh) was stimulated using 470 nm light pulses of 5ms duration and 8 Hz frequency. Inhibitory opsin (Jaws) was stimulated using 520 nm light pulses of 500 ms duration and 0.5 Hz frequency. The experiment followed a schedule of paired ON and OFF days after the end of training phase (DS + PS). The optical stimulation patch cord was plugged onto the ferrule during all experimental sessions (ON and OFF days) to habituate animals and control for latent experimental effects.

### *Ex vivo* patch-clamp recordings of VTA DA neurons

To verify the functional expression of the excitatory opsin CatCh and the inhibitory opsin Jaws, 10-12 week-old male DAT^iCRE^ mice were injected with the viruses described above. After 4 weeks, mice were deeply anesthetized with an intraperitoneal (IP) injection of a mix of ketamine/xylazine. Coronal midbrain sections (250 μm) were sliced using a Compresstome (VF-200; Precisionary Instruments) after intracardial perfusion of cold (4°C) sucrose-based artificial cerebrospinal fluid (SB-aCSF) containing (in mM): 125 NaCl, 2.5 KCl, 1.25 NaH_2_PO_4_, 5.9 MgCl_2_, 26 NaHCO_3_, 25 Sucrose, 2.5 Glucose, 1 Kynurenate (pH 7.2, 325 mOsm). After 10-60 min at 35°C for recovery, slices were transferred into oxygenated aCSF containing (in mM): 125 NaCl, 2.5 KCl, 1.25 NaH_2_PO_4_, 2 CaCl_2_, 1 MgCl_2_, 26 NaHCO_3_, 15 Sucrose, 10 Glucose (pH 7.2, 325 mOsm) at room temperature for the rest of the day and individually transferred to a recording chamber continuously perfused at 2 ml/min with oxygenated aCSF. Patch pipettes (4–8 MΩ) were pulled from thin wall borosilicate glass (G150TF-3, Warner Instruments) using a micropipette puller (P-87, Sutter Instruments, Novato, CA) and filled with a KGlu based intra-pipette solution containing (in mM): 116 K-gluconate, 10-20 HEPES, 0.5 EGTA, 6 KCl, 2 NaCl, 4 ATP, 0.3 GTP and 2 mg/mL biocytin (pH adjusted to 7.2). Transfected VTA DA cells were visualized using an upright microscope coupled with a Dodt contrast lens and illuminated with a white light source (Scientifica). To characterize CatCh expression, a 460 nm LED (CoolLED) was used both for visualizing YFP positive cells (using a bandpass filter cube) and for optical stimulation through the microscope (1s continuous for light-evoked current in voltage-clamp mode and 8 Hz with 5 ms/pulses to drive neuronal firing in current-clamp mode). Regarding Jaws expression, continuous photostimulation (20 s), with a 525 nm, pE-2, CoolLED, was used in current-clamp (−60 mV). Whole-cell recordings were performed using a patch-clamp amplifier (Axoclamp 200B, Molecular Devices) connected to a Digidata (1550 LowNoise acquisition system, Molecular Devices). Signals were low pass filtered (Bessel, 2 kHz) and collected at 10 kHz using the data acquisition software pClamp 10.5 (Molecular Devices). All the electrophysiological recordings were extracted using Clampfit (Molecular Devices) and analyzed with R.

### *In vivo* juxtacellular recordings of VTA DA neurons

Mice were deeply anaesthetized with chloral hydrate (8%), 400 mg/kg IP, supplemented as required to maintain optimal anesthesia throughout the experiment. The scalp was opened and a hole was drilled in the skull above the location of the VTA. Extracellular recording electrodes were constructed from 1.5 mm O.D. / 1.17 mm I.D. borosilicate glass tubings (Harvard Apparatus) using a vertical electrode puller (Narishige). Under microscopic control, the tip was broken to obtain a diameter of approximately 1 μm. The electrodes were filled with a 0.5% NaCl solution containing 1.5% of Neurobiotin tracer (AbCys) yielding impedances of 6-9 MΩ. Electrical signals were amplified by a high-impedance amplifier (Axon Instruments) and monitored through an audio monitor (A.M. Systems Inc.). The signal was digitized, sampled at 25 kHz and recorded using Spike2 software (Cambridge Electronic Design) for later analysis. The electrophysiological activity was sampled in the central region of the VTA (coordinates: between 3.1 to 4 mm posterior to bregma, 0.3 to 0.7 mm lateral to midline, and 4 to 4.8 mm below brain surface). Individual electrode tracks were separated from one another by at least 0.1 mm in the horizontal plane. Spontaneously active DA neurons were identified based on previously established electrophysiological criteria ^54,55^

### Fluorescence immunohistochemistry

After euthanasia, induced by IP injection of euthasol (0.1 mL per 30 g at 150 mg/kg) or by paraformaldehyde (PFA) intra-cardiac perfusion, brains were rapidly removed and fixed in 4% PFA. Following a period of fixation at 4°C, serial 60-μm sections were cut from the midbrain with a vibratome. Immunohistochemistry was performed as follows: free-floating VTA brain sections were incubated 1h at 4°C in a blocking solution of phosphate-buffered saline (PBS) containing 3% Bovine Serum Albumin (BSA, Sigma A4503) and 0.2% Triton X-100 and then incubated overnight at 4°C with a mouse anti-tyrosine hydroxylase antibody (TH, Sigma, T1299) at 1:200 dilution in PBS containing 1.5% BSA and 0.2% Triton X-100. The following day, sections were rinsed with PBS and then incubated for 3h at 22–25 °C with Cy3-conjugated anti-mouse (Jackson ImmunoResearch, 715-165-150) at 1:200 dilution in a solution of 1.5% BSA in PBS, respectively. After three rinses in PBS, slices were wet-mounted using Prolong Gold Antifade Reagent (Invitrogen, P36930). Microscopy was carried out with a fluorescent microscope Leica DMR, and images captured in grey level using MetaView software (Universal Imaging Corporation) and colored post-acquisition with ImageJ.

For the optogenetic experiments on DAT^iCRE^ mice, an immunohistochemical identification of the transfected neurons was performed as described above, with an addition of chicken anti-eYFP antibodies (Life technologies Molecular Probes, A-6455), at 1:500 dilution. A goat-anti-chicken AlexaFluor 488 secondary antibody (711-225-152, Jackson ImmunoResearch) at 1:500 dilution (Life Technologies) was then used in a solution of 1.5% BSA in PBS. Neurons co-labelled for TH and YFP in the VTA allowed to confirm their neurochemical phenotype and the transfection success.

### Intracranial self-stimulation (ICSS) bandit task

#### Behavioral set up

The ICSS bandit task took place in a circular open field with a diameter of 67 cm. Three explicit square-shaped marks (1×1 cm) were placed in the open field, forming an equilateral triangle (side = 35 cm). Entry in the circular zones (diameter = 6 cm) around each mark was associated with the delivery of a rewarding ICSS stimulation. Experiments were performed using a video camera, connected to a video-tracking system, out of sight of the experimenter. A LabVIEW (National Instruments) application precisely tracked and recorded the animal’s position with a camera (20 frames/s). When a mouse was detected in one of the circular rewarding zones, an electrical stimulator received a TTL signal from the software application and generated a 200 ms-train of 0.5-ms biphasic square waves pulsed at 100 Hz (20 pulses per train). ICSS intensity was adjusted, within a range of 20 to 200 μA, during training (see training settings) and then kept constant, so that mice would achieve between 50 and 150 visits per session (5min duration) for two successive sessions, and then kept constant for all the experiment. Mice with insufficient scores in the PS and DS (<40 visits despite increasing the maximum intensity of 200 μA) were excluded.

#### Training setting

The training consisted of two settings: the deterministic setting (DS) and the probabilistic setting (PS), both consisting of 10 daily sessions of 5 min. In the DS, all zones were associated with an ICSS delivery (P = 100%). However, two consecutive rewards could not be delivered on the same target, which motivates mice to alternate between targets. In the PS, the zones were associated with three different probabilities (P = 25%, P = 50%, P = 100%) to obtain an ICSS stimulation. The probabilities locations were pseudo-randomly assigned per mouse.

#### Data acquisition per experimental group

Different experimental groups underwent the ICSS bandit task. Firstly, locomotion and choice behavior of the mice, which had been implanted with osmotic mini-pumps (Sal = 23, Nic = 27), were analyzed and compared between the last two days of both training settings (days 9&10 (DS) + days 9&10 (PS)). For optogenetics experiments, the DAT^iCRE^ mice (n = 21) completed the training, followed by a schedule of 4 days of paired sessions with photo-stimulation (ON) alternated with days without photostimulation (OFF). The averages of the ON and OFF days were compared in a paired manner. The Ctl animals (n = 55) were obtained by pooling together mice implanted with a saline mini-pump (n = 23) and non-implanted mice (n = 32). Figure 1 used data from the non-implanted mice group. Figure 2,3,4 used the pooled Ctl group.

#### Behavioral measures

For all of those groups, the following measures were analyzed and compared in the PS, as well as in the DS for the Sal vs Nic experiment: i) number of visits, ii) time-to-goal, iii) choice repartition (proportion of visits p_25_, p_50_ and p_100_), iv) percentage of directional changes (n^th^ visit = n^th^ visit+2). Furthermore, the ICSS bandit task can be seen as a Markovian decision process. Every transition between zones can be considered as a binary choice between two probabilities, since the occupied zone cannot be reinforced twice in a row. The sequence of choices per session is summarized by the proportional result of the sum of three specific binary choices (or gambles, i.e., total visits zone 1/total visits zone 1+2). The three gambles (G) were named after the point on which the mouse is positioned at the time of the choice: G_25_ = 100 % vs 50 %, G_100_ = 50 % vs 25 % and G_50_ = 100 % vs 25 %. The target choice in these gambles reflects the balance between exploitative (choosing the most valuable option) and exploratory (choosing the least valuable option) choices. With a softmax based decision-making model fitted in the laboratory, we computed three parameters: the value sensitivity or inverse temperature (the power to discriminate between values in a binary choice), the uncertainty bonus (the preference for expected uncertainty, considering the reward variance of every option in a binary choice) and the motor cost to do a directional change (a decrease in the target value if it requires to go back to the previous target).

#### Modeling

Decision-making models determined the probability *P_i_* of choosing the next state i, as a function (the “choice rule”) of a “decision variable”. Because mice could not return to the same rewarding target, they had to choose between the two remaining ones. Accordingly, we modeled decisions between two alternatives labelled A and B and used a softmax choice rule defined by P_A =_1 / (1+e^−ß(vA-vB)^) where β is an inverse temperature parameter reflecting the sensitivity of choice to the difference between decision variables and *V_i_* the value of an option. The value *V* of an option is modelled as the expected (average) reward + expected uncertainty + U-turn cost ^16,30^. This compound value is then nested in the softmax choice rule, given a 6*3 matrix that described the probability of a choice between A, B and C (the three targets) depending on the two previous choices. As an example, in the probability to choose (A, B, C) after performing the sequence BA, the value is given by (0, p_b_ + φ∗p_b_*(1-p_b_)−κ, p_c_ + φ ∗ p_c_*(1-p_c_)) while after the sequence CA the value is given by (0, p_b_+φ∗p_b*_(1-p_b_), p_c_+φ∗p_c_*(1-p_c_) −κ) (same for AB, CB and AC, BC). The free parameters of the model (β, φ, κ) were fitted by maximizing the data likelihood. Given a sequence of choice c = c_1..T_, data likelihood is the product of their probability (given by Equation 1) ^56^. We used the *optim* function in R to perform the fits, with the constraints that β ∈]0,10], φ ∈]0,5] and κ ∈]0,5].

#### Statistical analysis

All statistical analyses were computed using R (The R Project, version 4.0.0) and Python with custom programs. Results were plotted as a mean ± s.e.m. The total number (n) of observations in each group and the statistics used are indicated in figure legends. Classical comparisons between means were performed using parametric tests (Student’s T-test, or ANOVA for comparing more than two groups) when parameters followed a normal distribution (Shapiro test P > 0.05), and non-parametric tests (here, Wilcoxon or Mann-Whitney) when the distribution was skewed. Multiple comparisons were Bonferroni corrected. Probability distributions were compared using the Kolmogorov–Smirnov (KS) test, and proportions were evaluated using a chi-squared test (χ^2^).

All statistical tests were two-sided except for the optogenetic experiment (Figure 5) where statistical tests were one-sided (we test hypotheses driven by nicotine effect and model). P > 0.05 was considered not to be statistically significant. For archetypal analysis, all computations and graphics have been done using the statistical software R and the archetype package (version 2.2-0.1). Briefly, given an n × m matrix representing a multivariate data set with n observations (n = number of animals) and m attributes (here *m* = 7, consisting of the directional changes, the target distributions (3 values) and the three gambles (see data Figure 1 C-E)), the archetypal analysis finds the matrix *Z* of *k m*-dimensional archetypes (*k* is the number of archetypes). *Z* is obtained by minimizing || *X-α Z^T^* ||_2_, with α the coefficients of the archetypes *(α_i,1..k_ ≥0* and *∑ α_i,1..k_ = 1*), and ||.||_2_ a matrix norm. The archetype is also a convex combination of the data points *Z*=*X^T^δ,* with *δ* ≥ 0 and their sum must be 1 ^57^. The α-coefficient depicts the relative archetypal composition of a given observation. For *k =* 3 archetypes and an observation i, *α*_i,1_, *α*_i,2_, *α*_i,3_ ≥0 and *α*_i,1_ +α_i,2_ + *α*_i,3_ = 1. A ternary plot can then be used to visualize data. (*α*_i,1_, *α*_i,2_, *α*_i,2_) are used to assign individual behavior to its nearest archetype (i.e, k max(*α*_i,1_, *α*_i,2_, *α*_i,3_)). *α*_i,j_ are also used as variable to estimate population archetypal composition. For figure 4E, archetypal composition (*0≤α_i,j_ ≤1)* was binned into five intervals. Pure archetype corresponds to 1, the archetypal composition decreases linearly with increasing distance from the archetype, 0 correspond to points on the opposite side.

---

**Supplementary Figure 1:**
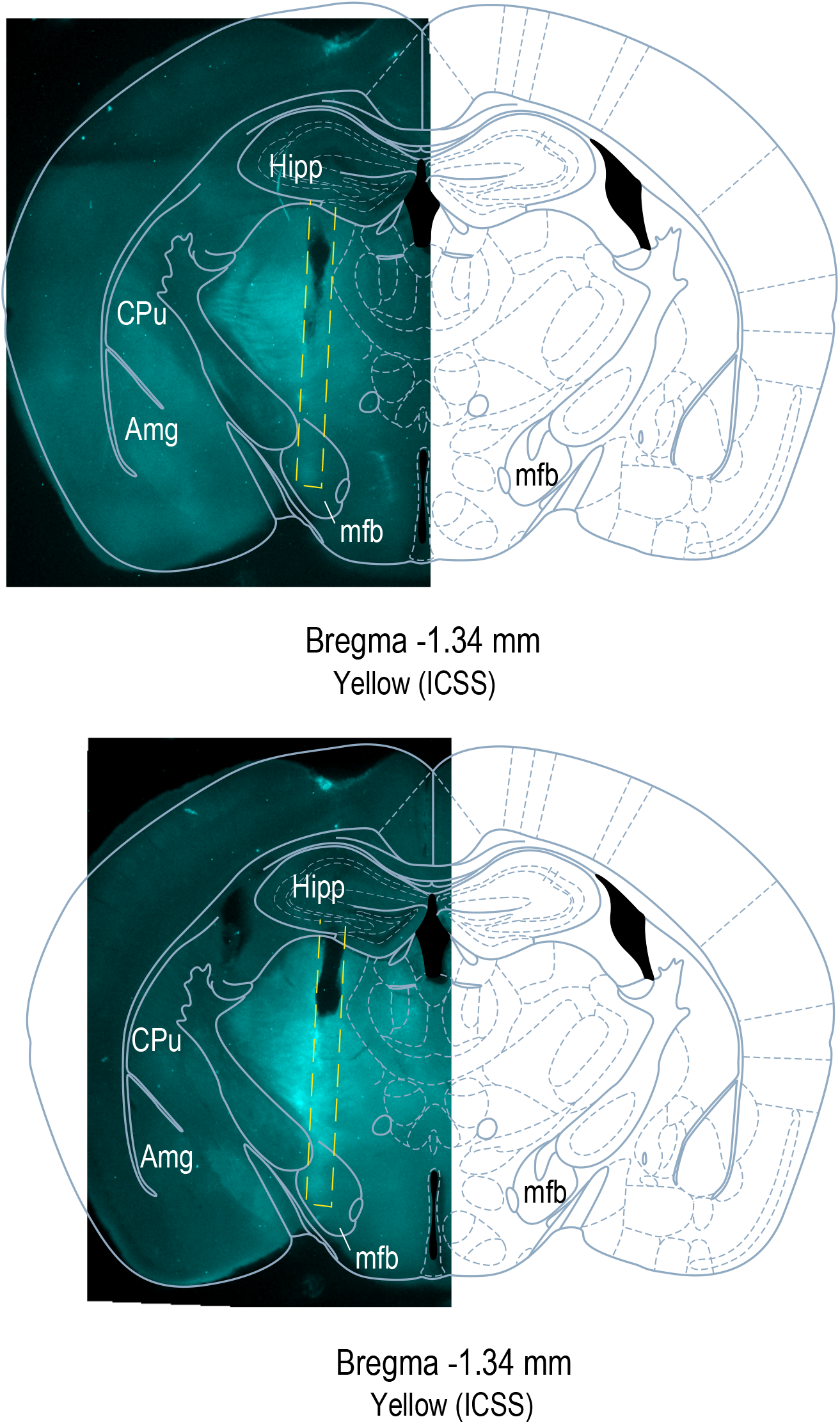
Stimulating electrode implantation: Representative examples of unilateral MFB implantations in two different brains. *Post-hoc* verification of the ICSS track is represented in dotted yellow line.

**Supplementary Figure 2:**
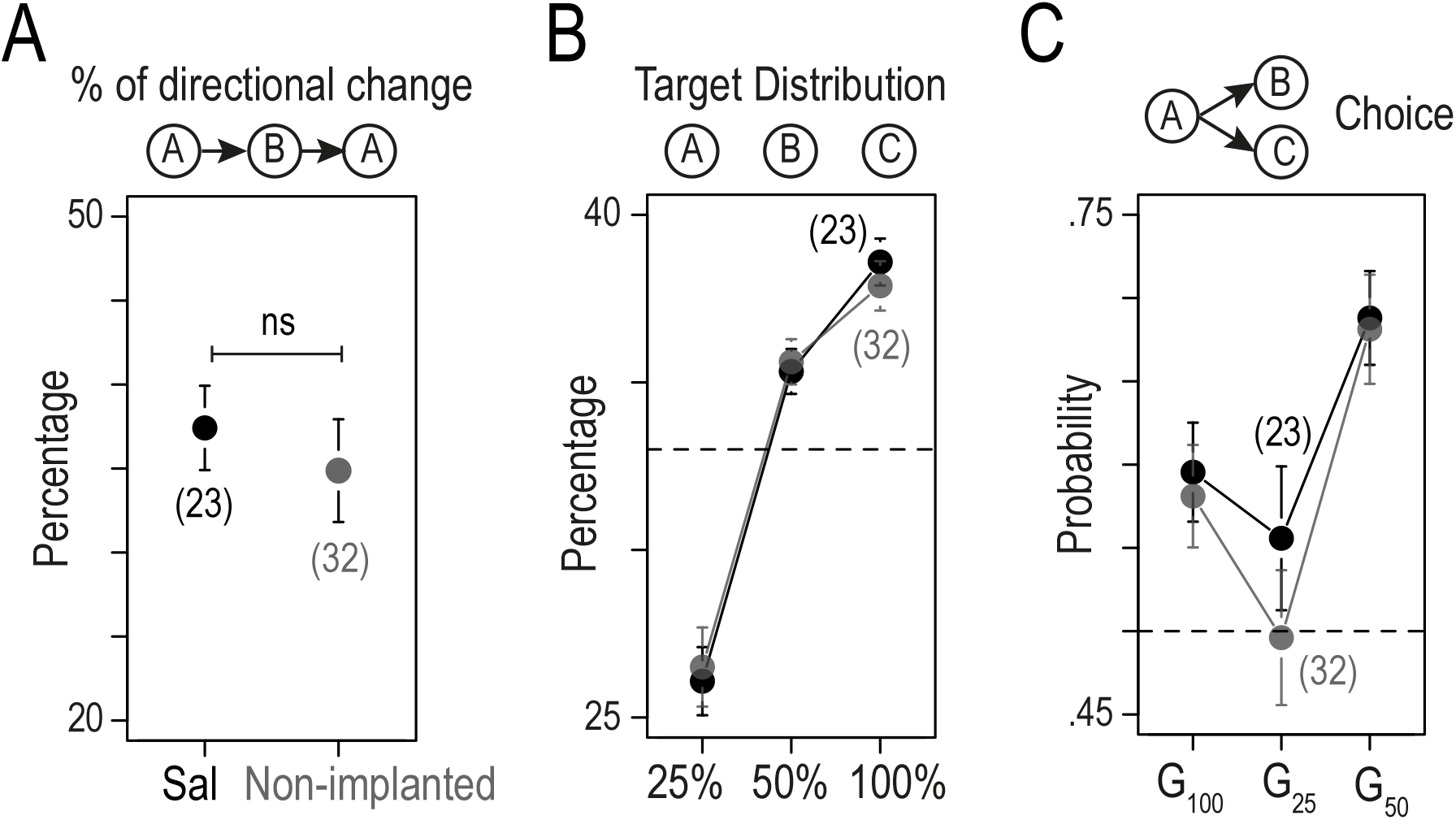
No behavioral difference between mice implanted with an osmotic minipump filled with saline (Sal) and non-implanted mice. (A) Comparison of the percentage of directional change in Sal (black) and non-implanted (grey) mice (Wilcoxon signed rank test, p >0.05). (B) Repartition of the visits to the three targets in the DS (Wilcoxon Test p >0.05 for the three comparisons). (C) Probability to choose the alternative with the highest probability of reward for the three possible gambles: G_100_; G_25_ and G_50_ (Wilcoxon Test p >0.05 for the three comparisons)

**Supplementary Figure 3:**
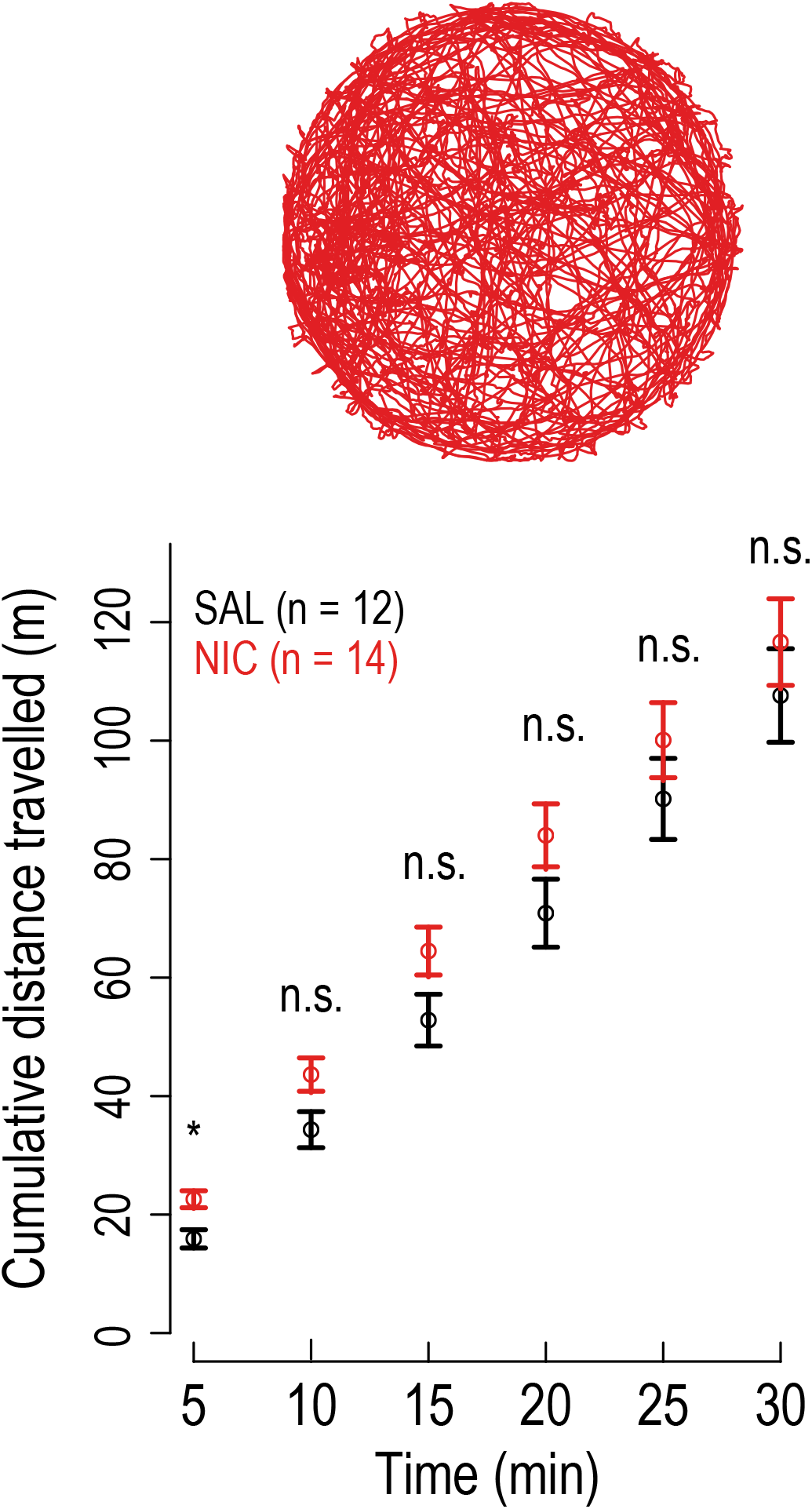
Nicotine-treated mice show increased locomotion for the first five minutes in an open field. Top, trajectory in an open field (duration 30 minutes) of a mouse treated for 24 days with nicotine (10 mg/kg/day). Bottom, cumulative distance travelled in meters measured every 5 minutes during a 30 min-OF exploration. Nic mice (n = 14) showed a greater distance travelled during the first 5 minutes only (t-test, *t* = −2.4154, *df* = 22.074, *p* = 0.02444), compared to saline-exposed mice (Sal, n = 12). The total distance travelled after 30 minutes was not significantly different between the two groups (p > 0.05).

**Supplementary Figure 4:**
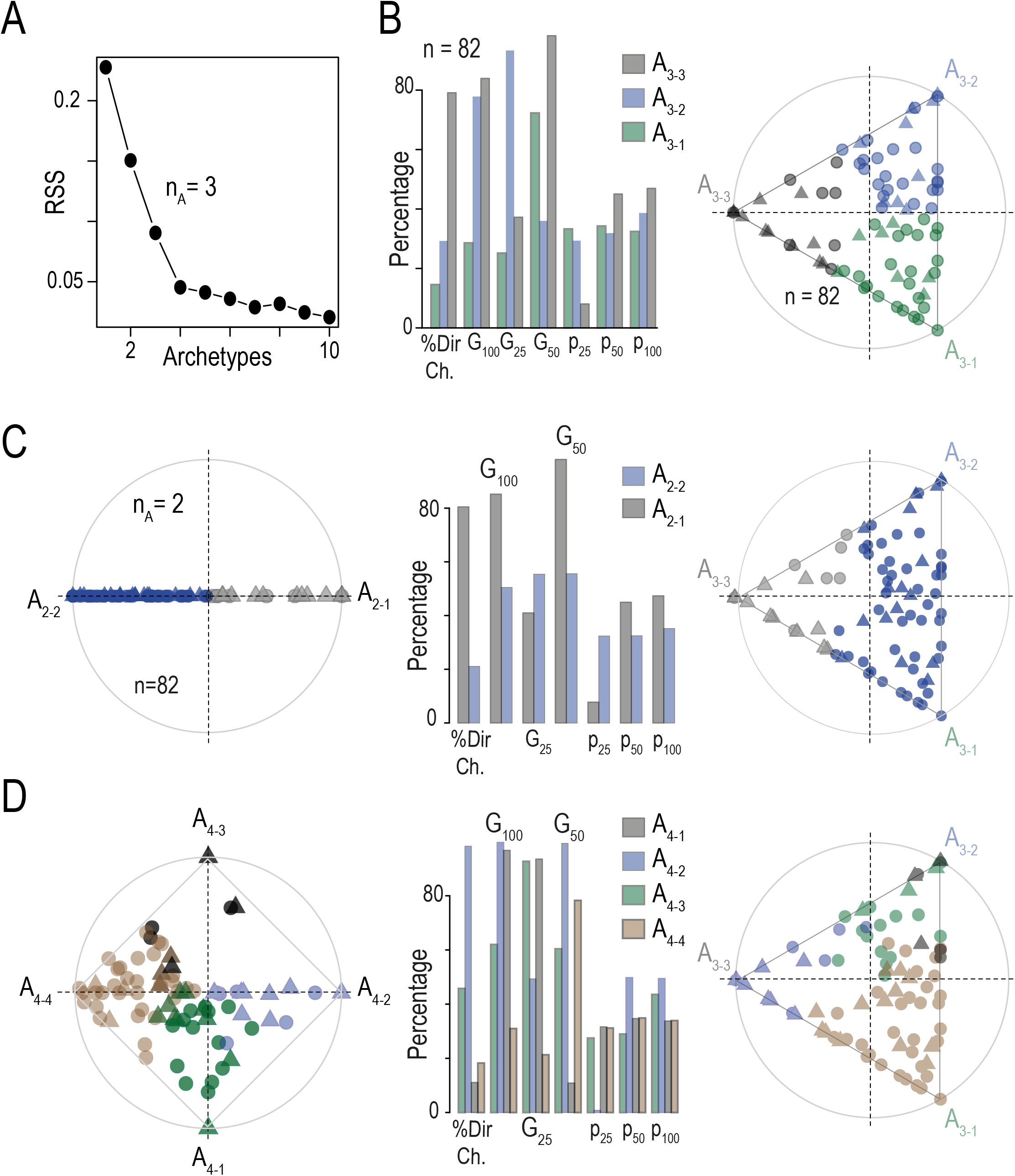
Archetypal analyses with 2 to 4 apices: Archetypal analysis of the choice strategies based on a 7-dimensional data space: % of directional change, Gambles G_100_ (choice 50% over 25%), G_25_ (100% over 50%) and G_50_ (100% over 25%), and probabilities of choosing each point (P_25_, P_50_, and P_100_). Analyses were performed on n = 82 mice (pooled Ctl and Nic mice). (A) Residual sum of squares for a number or archetypes n_A_ = 1 to 10. Error reduction between n_A_ = 4 and 10 is marginal. (B) Plot of the archetypal solutions for n_A_ = 3. *Left*: Percentile plot of the value of the 7 basic variables used in this analysis for the three archetypes, here labelled A_3-_ 1 to A_3-3_. A_3-1_, A_3-2_ and A_3-3_ correspond to the GO, Des and Asc archetypes of Figure 3D, but are unlabeled here for comparison purposes with n_A_ = 2 and 4. *Right*: Visualization of the α coefficients using a ternary plot, in which the three apices represent the three archetypes. Each point shows the projection of each individual (n=82). Points are color-coded according to their proximity to the archetypes. (C) Plot of the archetypal solutions A_2-1_ and A_2-2_ for n_A_ = 2. From left to right: Visualization of the α coefficients in a binary plot, percentile plot and ternary plot (same as in B, with points color-coded according to their proximity to the two archetypes A_2-1_ and A_2-2_). (D) Same as C for n_A_ = 4 (A_4-1_ to A_4-4_). Note that the A_3-3_ (a.k.a. GO) archetype is present when both n_A_ = 2 (the A_2-1_ archetype) and n_A_ = 4 (A_4-2_).

**Supplementary Figure 5:**
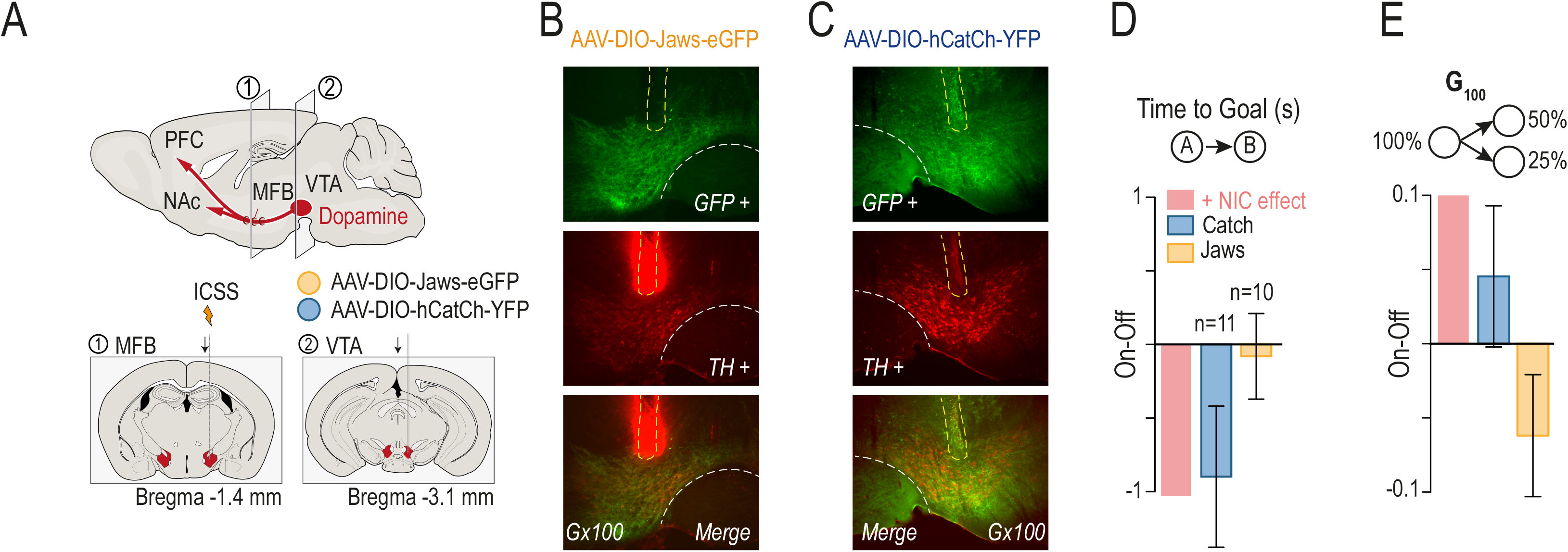
Optogenetic manipulation of choice behaviors. (A) DAT-Cre mice were implanted unilaterally with bipolar stimulating electrodes for ICSS in the medial forebrain bundle (1) and transduced with either an AAV-DIO-Jaws-eGFP or an AAV-DIO-Catch-YFP in the VTA (2). (B) Representative immunohistochemical verification of CatCh-YFP expression selectively in DA neurons of the VTA (anti-TH in red, and -GFP in green; merge on top). *Post-hoc* verification of the unilateral fiber implantation is represented in dotted yellow lines. (C) Representative immunohistochemical verification of Jaws-eGFP expression selectively in DA neurons of the VTA (anti-TH in red, and -GFP in green; merge on top). *Post-hoc* verification of the unilateral fiber implantation is represented in dotted yellow lines. (D-E) Net effect of light stimulation on (D) the time to goal and (E) the percent of choice toward P_50_ in gamble G_100_. Data from OFF sessions were subtracted from data from ON sessions. In red, the effect of nicotine (NIC) is represented for both parameters, as a comparison factor. (Asterisk: Comparison with a true mean of 0 and paired comparison (ON – OFF) (Student’s t-test or Wilcoxon test, unilateral testing).

## References

1. Fowler, C. D., Arends, M. A. & Kenny, P. J. Subtypes of nicotinic acetylcholine receptors in nicotine reward, dependence, and withdrawal: evidence from genetically modified mice. Behav Pharmacol 19, 461–484 (2008).

2. Marti, F. et al. Smoke extracts and nicotine, but not tobacco extracts, potentiate firing and burst activity of ventral tegmental area dopaminergic neurons in mice. Neuropsychopharmacology 36, 2244–2257 (2011).

3. Stolerman, I. P. & Jarvis, M. J. The scientific case that nicotine is addictive. Psychopharmacology 117, 2–10– discussion 14–20 (1995).

4. Lüscher, C. & Malenka, R. C. Drug-evoked synaptic plasticity in addiction: from molecular changes to circuit remodeling. Neuron 69, 650–663 (2011).

5. Association, A. P. Diagnostic and statistical manual of mental disorders (DSM-5®). (2013).

6. Naudé, J., Dongelmans, M. & Faure, P. Nicotinic alteration of decision-making. Neuropharmacology 96, 244–254 (2015).

7. Addicott, M. A., Pearson, J. M., Sweitzer, M. M., Barack, D. L. & Platt, M. L. A Primer on Foraging and the Explore/Exploit Trade-Off for Psychiatry Research. Neuropsychopharmacology 42, 1931–1939 (2017).

8. Locey, M. L. & Dallery, J. Isolating behavioral mechanisms of intertemporal choice: nicotine effects on delay discounting and amount sensitivity. Journal of the experimental analysis of behavior 91, 213–223 (2009).

9. Viñals, X. et al. Overexpression of α3/α5/β4 nicotinic receptor subunits modifies impulsive-like behavior. Drug Alcohol Depend 122, 247–252 (2012).

10. Addicott, M. A., Pearson, J. M., Froeliger, B., Platt, M. L. & McClernon, F. J. Smoking automaticity and tolerance moderate brain activation during explore-exploit behavior. Psychiatry Research 224, 254–261 (2014).

11. Addicott, M. A., Pearson, J. M., Wilson, J., Platt, M. L. & McClernon, F. J. Smoking and the bandit: A preliminary study of smoker and nonsmoker differences in exploratory behavior measured with a multiarmed bandit task. Experimental and Clinical Psychopharmacology 21, 66–73 (2013).

12. Levine, A. et al. Molecular mechanism for a gateway drug: epigenetic changes initiated by nicotine prime gene expression by cocaine. Sci Transl Med 3, 107ra109 (2011).

13. Cohen, J. D., McClure, S. M. & Yu, A. J. Should I stay or should I go? How the human brain manages the trade-off between exploitation and exploration. Philos Trans R Soc Lond, B, Biol Sci 362, 933–942 (2007).

14. Wyart, V., Sciences, E. K. C. O. I. B.2016. Choice variability and suboptimality in uncertain environments. Personality and Individual Differences 11, 109–115 (2016).

15. Wilson, R. C., Geana, A., White, J. M., Ludvig, E. A. & Cohen, J. D. Humans use directed and random exploration to solve the explore-exploit dilemma. J Exp Psychol Gen 143, 2074–2081 (2014).

16. Belkaid, M. et al. Mice adaptively generate choice variability in a deterministic task. Communications Biology 3, 1–9 (2020).

17. Berlyne, D. E. Curiosity and exploration. Science 153, 25–33 (1966).

18. Schultz, W. Multiple dopamine functions at different time courses. Annu Rev Neurosci 30, 259–288 (2007).

19. Redish, A. D., Jensen, S. & Johnson, A. A unified framework for addiction: Vulnerabilities in the decision process. The Behavioral and brain sciences 31, 415–37; discussion 437–87 (2008).

20. Lüscher, C., Robbins, T. W. & Everitt, B. J. The transition to compulsion in addiction. Nat Rev Neurosci 21, 1–17 (2020).

21. Kalivas, P. W. & Volkow, N. D. The neural basis of addiction: a pathology of motivation and choice. 162, 1403–1413 (2005).

22. Mizumori, S. J. Y. & Jo, Y. S. Homeostatic regulation of memory systems and adaptive decisions. Hippocampus 23, 1103–1124 (2013).

23. Cagniard, B. et al. Dopamine scales performance in the absence of new learning. Neuron 51, 541–547 (2006).

24. Westbrook, A. & Braver, T. S. Dopamine Does Double Duty in Motivating Cognitive Effort. Neuron 89, 695–710 (2016).

25. Niv, Y., Daw, N. D., Joel, D. & Dayan, P. Tonic dopamine: opportunity costs and the control of response vigor. Psychopharmacology 191, 507–520 (2007).

26. Frank, M. J., Doll, B. B., Oas-Terpstra, J. & Moreno, F. Prefrontal and striatal dopaminergic genes predict individual differences in exploration and exploitation. Nat Neurosci 12, 1062–1068 (2009).

27. Humphries, M. D., Khamassi, M. & Gurney, K. Dopaminergic Control of the Exploration-Exploitation Trade-Off via the Basal Ganglia. Frontiers in neuroscience 6, 9 (2012).

28. Beeler, J. A., Daw, N., Frazier, C. R. M. & Zhuang, X. Tonic dopamine modulates exploitation of reward learning. Front. Behav. Neurosci. 4, 170 (2010).

29. Cinotti, F. et al. Dopamine blockade impairs the exploration-exploitation trade-off in rats. Sci. Rep. 9, 6770 (2019).

30. Naudé, J. et al. Nicotinic receptors in the ventral tegmental area promote uncertainty-seeking. Nat Neurosci 19, 471–478 (2016).

31. Faure, P., Tolu, S., Valverde, S. & Naudé, J. Role of nicotinic acetylcholine receptors in regulating dopamine neuron activity. Neuroscience 282C, 86–100 (2014).

32. Epping-Jordan, M. P., Watkins, S. S., Koob, G. F. & Markou, A. Dramatic decreases in brain reward function during nicotine withdrawal. Nature 393, 76–79 (1998).

33. Morel, C. et al. Nicotinic receptors mediate stress-nicotine detrimental interplay via dopamine cells’ activity. Mol Psychiatry 23, 1597–1605 (2017).

34. Tolu, S. et al. Nicotine enhances alcohol intake and dopaminergic responses through β2* and β4* nicotinic acetylcholine receptors. Sci. Rep. 7, 45116 (2017).

35. Cutler, A. & Breiman, L. Archetypal Analysis. Technometrics 36, 338–347 (1994).

36. Hart, Y. et al. Inferring biological tasks using Pareto analysis of high-dimensional data. Nat Meth 12, 233–5– 3 p following 235 (2015).

37. Belkaid, M. et al. Mice adaptively generate choice variability in a deterministic task - behavioral data. (2019). doi:10.5281/zenodo.3576423

38. Besson, M. et al. Long-term effects of chronic nicotine exposure on brain nicotinic receptors. Proc Natl Acad Sci USA 104, 8155–8160 (2007).

39. Kleinlogel, S. et al. Ultra light-sensitive and fast neuronal activation with the Ca^2^+-permeable channelrhodopsin CatCh. 14, 513–518 (2011).

40. Chuong, A. S. et al. Noninvasive optical inhibition with a red-shifted microbial rhodopsin. 17, 1123–1129 (Nature Publishing Group, 2014).

41. Heilbronner, S. R. & Hayden, B. Y. Contextual factors explain risk-seeking preferences in rhesus monkeys. Frontiers in neuroscience 7, 7 (2013).

42. Hernandez, G., Trujillo-Pisanty, I., Cossette, M.-P., Conover, K. & Shizgal, P. Role of dopamine tone in the pursuit of brain stimulation reward. Journal of Neuroscience 32, 11032–11041 (2012).

43. Kenny, P. J. & Markou, A. Nicotine self-administration acutely activates brain reward systems and induces a long-lasting increase in reward sensitivity. Neuropsychopharmacology 31, 1203–1211 (2006).

44. Juarez, B. et al. Midbrain circuit regulation of individual alcohol drinking behaviors in mice. Nature Communications 8, 2220 (2017).

45. Stern, S., Kirst, C. & Bargmann, C. I. Neuromodulatory Control of Long-Term Behavioral Patterns and Individuality across Development. Cell 171, 1–25 (2017).

46. MacDonald, S. W. S., Nyberg, L. & Bäckman, L. Intra-individual variability in behavior: links to brain structure, neurotransmission and neuronal activity. TINS 29, 474–480 (2006).

47. Torquet, N. et al. Social interactions impact on the dopaminergic system and drive individuality. Nature Communications 9, 3081 (2018).

48. Smillie, L. D. & Wacker, J. Dopaminergic foundations of personality and individual differences. Front. Hum. Neurosci. 8, 874 (2014).

49. DeYoung, C. G. Personality Neuroscience and the Biology of Traits. Social and Personality Psychology Compass 4, 1165–1180 (2010).

50. Palmatier, M. I. et al. Dissociating the primary reinforcing and reinforcement-enhancing effects of nicotine using a rat self-administration paradigm with concurrently available drug and environmental reinforcers. Psychopharmacology 184, 391–400 (2006).

51. Chiu, P. H., Lohrenz, T. M. & Montague, P. R. Smokers’ brains compute, but ignore, a fictive error signal in a sequential investment task. Nat Neurosci 11, 514–520 (2008).

52. Addicott, M. A., Pearson, J. M., Kaiser, N., Platt, M. L. & McClernon, F. J. Suboptimal foraging behavior: a new perspective on gambling. Behav. Neurosci. 129, 656–665 (2015).

53. McGrath, D. S. & Barrett, S. P. The comorbidity of tobacco smoking and gambling: a review of the literature. Drug Alcohol Rev 28, 676–681 (2009).

54. Exley, R. et al. Distinct contributions of nicotinic acetylcholine receptor subunit alpha4 and subunit alpha6 to the reinforcing effects of nicotine. Proc Natl Acad Sci USA 108, 7577–7582 (2011).

55. Ungless, M. A. & Grace, A. A. Are you or aren’t you? Challenges associated with physiologically identifying dopamine neurons. TINS 35, 422–430 (2012).

56. Daw, N. D. in Decision Making, Affect, and Learning 3–38 (Oxford University Press). doi:10.1093/acprof:oso/9780199600434.003.0001

57. Eugster, M. J. A. & Leisch, F. From Spider-Man to Hero - Archetypal Analysis in R. Journal of Statistical Software 30, 1–23 (2009).

